# Kinetic analysis of ASIC1a delineates conformational signaling from proton-sensing domains to the channel gate

**DOI:** 10.1101/2021.01.18.427137

**Authors:** Sabrina Vullo, Nicolas Ambrosio, Jan P. Kucera, Olivier Bignucolo, Stephan Kellenberger

**Author notes:** To whom correspondence should be addressed: Stephan Kellenberger, Department of biomedical Sciences, University of Lausanne, Rue du Bugnon 27, CH-1011 Lausanne, Switzerland.

## Abstract

Acid-sensing ion channels (ASICs) are neuronal Na^+^ channels that are activated by a drop in pH. Their established physiological and pathological roles, involving fear behaviors, learning, pain sensation and neurodegeneration after stroke, make them promising targets for future drugs. Currently, the ASIC activation mechanism is not understood. Here we used voltage-clamp fluorometry (VCF) combined with fluorophore-quencher pairing to determine the kinetics and direction of movements. Molecular dynamics simulations were used to further evaluate VCF-predicted movements. We show that conformational changes with the speed of channel activation occur close to the gate and in more distant extracellular sites, where they may be driven by local protonation events. Further, we provide evidence for fast conformational changes in a pathway linking protonation sites to the channel pore, in which an extracellular interdomain loop interacts via aromatic residue interactions with the upper end of a transmembrane helix and would thereby open the gate.

## Introduction

This study investigates the activation mechanism of acid-sensing ion channels (ASICs), a family of H^+^-gated Na^+^ channels of the nervous system (Waldmann et al., 1997, Wemmie et al., 2013, Kellenberger and Schild, 2015, Yang and Palmer, 2014). ASIC activation is linked to physiological and pathological processes such as learning and pain sensation, neurodegeneration after ischemic stroke, fear and anxiety (rev. in (Wemmie et al., 2013, Kellenberger and Schild, 2015)). ASICs respond to extracellular acidification with a transient current, because after opening, they enter a non-conducting desensitized state (Waldmann et al., 1997, Grunder and Pusch, 2015). High-resolution structures of chicken ASIC1a (cASIC1a) in the closed (Yoder and Gouaux, 2020, Yoder et al., 2018) (and (Sun et al., 2020), human ASIC1a), toxin-opened (Baconguis et al., 2014; Baconguis and Gouaux, 2012; Dawson et al., 2012), and desensitized conformation (Gonzales et al., 2009; Jasti et al., 2007) are available. Functional ASICs are trimers (Bartoi et al., 2014). Each ASIC subunit consists of short intracellular *N*-and *C*-terminal ends, two transmembrane domains TM1 and TM2, and a large extracellular region, with the shape of a hand, organized in defined domains that have been named palm, knuckle, β-ball, thumb and finger (Figure 1A).

**Figure 1.**
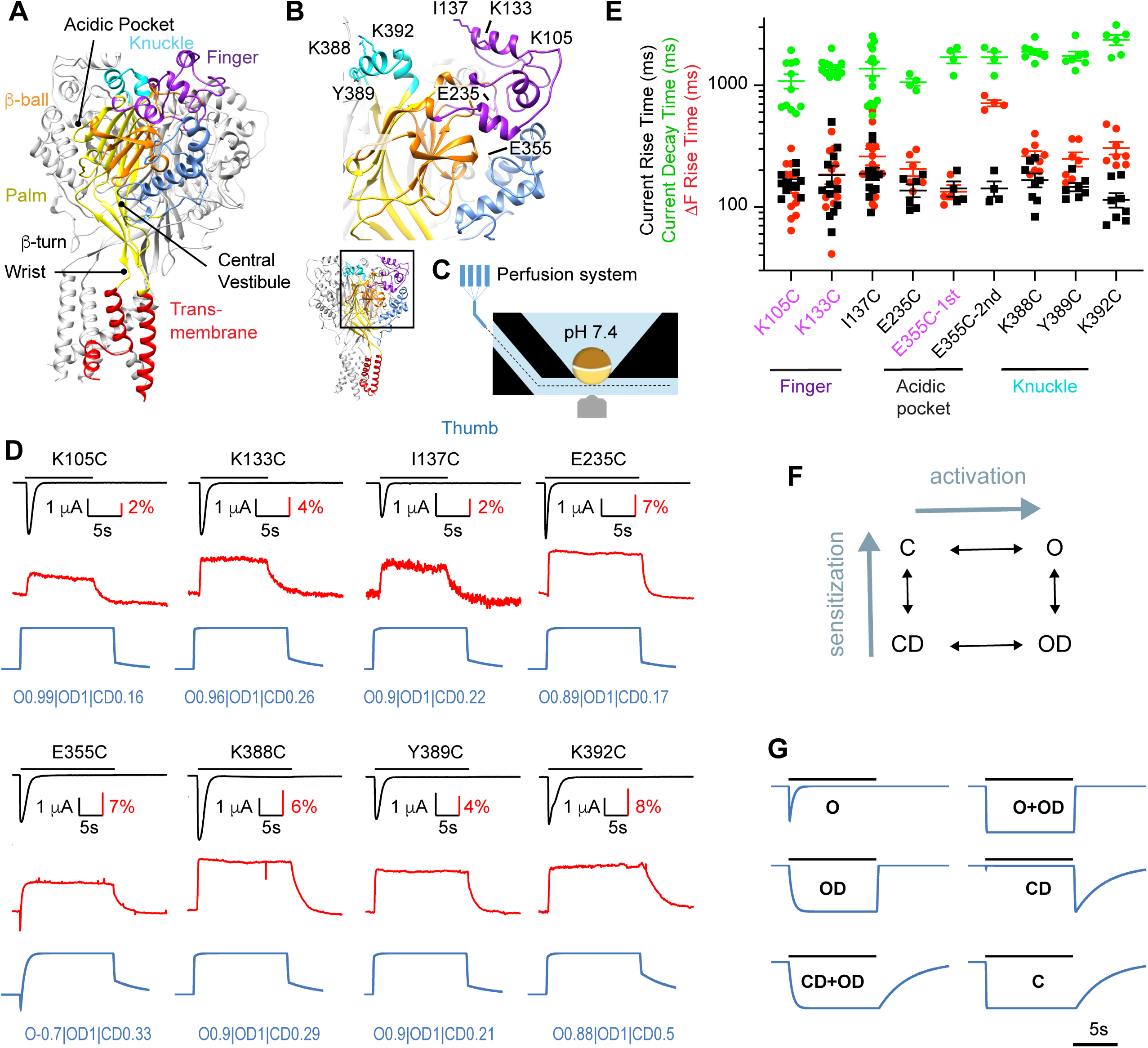
Rapid ΔF signals of mutants distant from the pore. (A) Structural image of a trimeric ASIC1a channel with the domains colored and named, based on the Mit Toxin-opened structure (PDB identifier 4NTW, (Baconguis et al., 2014)); the acidic pocket, central vestibule, β-turn and wrist are indicated. (**B**) Detailed structural view with indication of residues studied in Figure 1. (**C**) Schematic view of the oocyte chamber used for the kinetic measurements of currents and fluorescence signals (ΔF). **(D)** Current traces (black), fluorescence signals (red) and simulated fluorescence traces (blue) are shown. The conditioning pH between stimulations was pH7.4; the stimulation pH6 was applied as indicated by the horizontal black bars. Simulated traces were generated by the kinetic model (*Materials and Methods)*, using the parameters indicated in blue (Table S2). **(E)** Kinetics of current appearance and decay, and of ΔF onset, measured as rise time (RT) or decay time for an activation by pH6.0 (n=4-14). An analysis was carried out to determine if the ΔF onset kinetics were correlated with current appearance or decay (*Materials and Methods*). In mutants labeled in purple, the ΔF onset kinetics were correlated with current appearance. For E355C, ΔF onset kinetics are reported for both components (“1^st^” and “2^nd^”). **(F)** Representation of the kinetic model used for the simulation of fluorescence traces; C, closed; O, open; OD, open-desensitized; CD, closed-desensitized. **(G)** Model-generated fluorescence with proportionality factor =-1 for the state(s) indicated with the traces; for C, this factor was +1. Conditioning pH was 7.4; it was changed to pH6.0 for a duration of 10s. The fall time of the pH change (speed of solution change) was set to 200 ms. Source data are provided in the file Figure1-Source_Data.

Each ASIC channel contains three “acidic pockets” - regions with a high density of acidic residues - which are enclosed by the thumb, finger and β-ball of one, and the palm of a neighboring subunit. The lower palm domains enclose the central vestibule. The wrist links the extracellular channel parts to the transmembrane segments. The acidic pocket, the palm and the wrist are pH-sensing regions that are potentially involved in ASIC activation (Paukert et al., 2008, Krauson et al., 2013, Liechti et al., 2010, Vullo et al., 2017, Schuhmacher et al., 2015). It is expected that protonation would locally induce structural rearrangements, and these conformational changes would be transmitted to the gate. It may also be possible that the opening of the ASIC1a gate is controlled by protonation events in the wrist alone, since combined mutation of two putative pH-sensing His residues in the wrist suppressed ASIC1a activation, while maintaining cell surface expression intact (Paukert et al., 2008).

By using voltage-clamp fluorometry (VCF), we show here that conformational changes with the kinetics of ASIC activation occur in the wrist and in several distal extracellular parts, consistent with pH sensing in different channel regions. To investigate possible pathways transmitting conformational changes from distant sites to the channel gate, we determined the kinetics and direction of conformational changes in the palm and the adjacent palm-thumb loops, using VCF combined with the introduction of fluorescence quencher groups. The fluorescence signals were analyzed with a kinetic model, and the predicted distance changes were further evaluated with Molecular Dynamics simulations. This analysis allowed us to propose sequences of conformational rearrangements in ASIC domains in the proximity of the β1-β2 linker during activation and desensitization.

## Results

### Fast conformational changes in ASIC domains that are distant from the pore

The aim of a first set of experiments was to compare the kinetics of conformational changes in different parts of ASIC1a. To this end, VCF was used, which measures simultaneously with the channel current, changes in fluorescence (ΔF) of a strategically placed fluorophore, as a readout of changes in its environment. ASIC1a constructs containing engineered Cys residues for the docking of maleimide derivatives of fluorophores were expressed in *Xenopus laevis* oocytes, and oocytes were exposed to maleimide derivatives of AlexaFluor488 or CF488A prior to the measurement. The ΔF signals detected in labeled mutants are due to fluorophores attached to the engineered Cys residues, since exposure of wild type (WT) ASIC1a to the maleimide derivatives of these two fluorophores does not lead to fluorescence changes upon acidification (Bonifacio et al., 2014).

In previous studies we had observed rapid ΔF signals with mutations located distantly from the channel gate (Bonifacio et al., 2014, Gwiazda et al., 2015, Vullo et al., 2017). For an accurate comparison of the kinetics of these mutants (Figure 1B), they were measured here in a measuring chamber in which the current and ΔF signal are measured from approximately the same oocyte surface ((Figure 1C, (Vullo et al., 2017)). The tested mutants produced transient currents upon extracellular acidification (black traces in Figure 1D). The ΔF signals of these mutants (red traces) were sustained; only E355C showed an additional first transient component. Figure 1E presents for acidification to pH6 the rise time (time to pass from 10% to 90% of the maximal amplitude, RT) of the ΔF onset (red symbols) and of the current appearance (black), and the decay time (90 to 10%) of the current desensitization (green). The kinetic analysis illustrates that for most of these mutants, the ΔF onset kinetics are similar to those of current appearance (Figure 1E). To determine how closely the conformational changes underlying the ΔF signals were associated with a functional transition, a correlation analysis was done for each mutant (*Materials and Methods)*. This indicated a correlation to channel activation for K105C, K133C and E355C (purple label).

The results obtained at pH6.5 and 5.5 (Figure 1-figure supplement 1) were comparable with those at pH6.0 shown here.

The ΔF signals indicate that the environment of the docked fluorophores changes, thus that the acidification induces conformational changes. The ΔF may be due to changes in solvent exposure of the fluorophore (Gandhi and Olcese, 2008, Cha and Bezanilla, 1997, Mannuzzu et al., 1996), or to changes of the exposure to nearby quenching groups (Pantazis and Olcese, 2012, Vullo et al., 2017). It is expected that conformational transitions that are not directly associated with a defined functional state also influence the fluorescence signal. As an additional strategy for testing the association of ΔF signals with specific transitions, ΔF traces were compared to traces generated by a kinetic model under the assumption that each functional state contributes with a proportionality factor that lies between −1 and +1 to the measured fluorescence. To this end, a kinetic ASIC1a model recently developed in our laboratory was used (Alijevic et al., 2020), which is based on the Hodgkin-Huxley formalism containing an activation and a sensitization gate. The model contains four states, an open (O), a closed (C) and two desensitized states, closed-desensitized (CD) and open-desensitized (OD; Figure 1F). This model reproduces the acid-induced ASIC1a activity well (Alijevic et al., 2020). Figure 1G shows ΔF traces generated by this model for a pH change from 7.4 to 6.0, under the assumption that in each case one single functional state, or a pair, O+OD (corresponding to the activation gate) and CD+OD (sensitization gate) contributes to the signal. The ΔF onset is most rapid for the fluorescence associated with the O or O+OD states. The CD state is associated with a substantial ΔF only at the return to pH7.4 (Figure 1G). At the end of the acidic pulse when the pH is changed back to 7.4, the return to the initial fluorescence level is slow if the fluorescence is associated with the CD state, consistent with the slow recovery from desensitization at pH7.4 (Alijevic et al., 2020, Rook et al., 2020).

By attributing proportionality factors between −1 and +1 to the 4 states of the model, based on the shape and the kinetics of experimental ΔF traces (*Materials and Methods,* experimental values of the ΔF kinetics at the return to the conditioning pH are provided in Supplementary Table S1), ΔF signals can be generated that closely resemble the measured ΔF signals, as illustrated by the blue traces shown in Figure 1D, where the proportionality factors (Supplementary Table S2) are indicated for each mutant. This model suggests high association of the ΔF signal with the O and OD state, and lower association with the CD state.

### Fast conformational changes in the wrist have the same timing as channel opening

It was then tested whether the ΔF kinetics were different if the fluorophores were placed closer to the channel gate. Mutations were therefore introduced in the wrist (Figure 2A). In contrast to the mutants at distant positions (Figure 1), the ΔF traces of these mutants had two components, a first transient and a second sustained part (Figure 2B). For transient ΔF components, the RT of the onset (“on”) and of the decay of the signal (“off”) were determined. The analysis shows that the kinetics of the fast ΔF component of all mutants were equal to or faster than current appearance (Figure 2C-D, *Methods*; pH6.5 and 5.5 data in Figure 2-figure supplement 1). The traces were best modeled with highest proportionality factors for the open state, highlighting the association of the ΔF of these mutants with channel opening.

**Figure 2.**
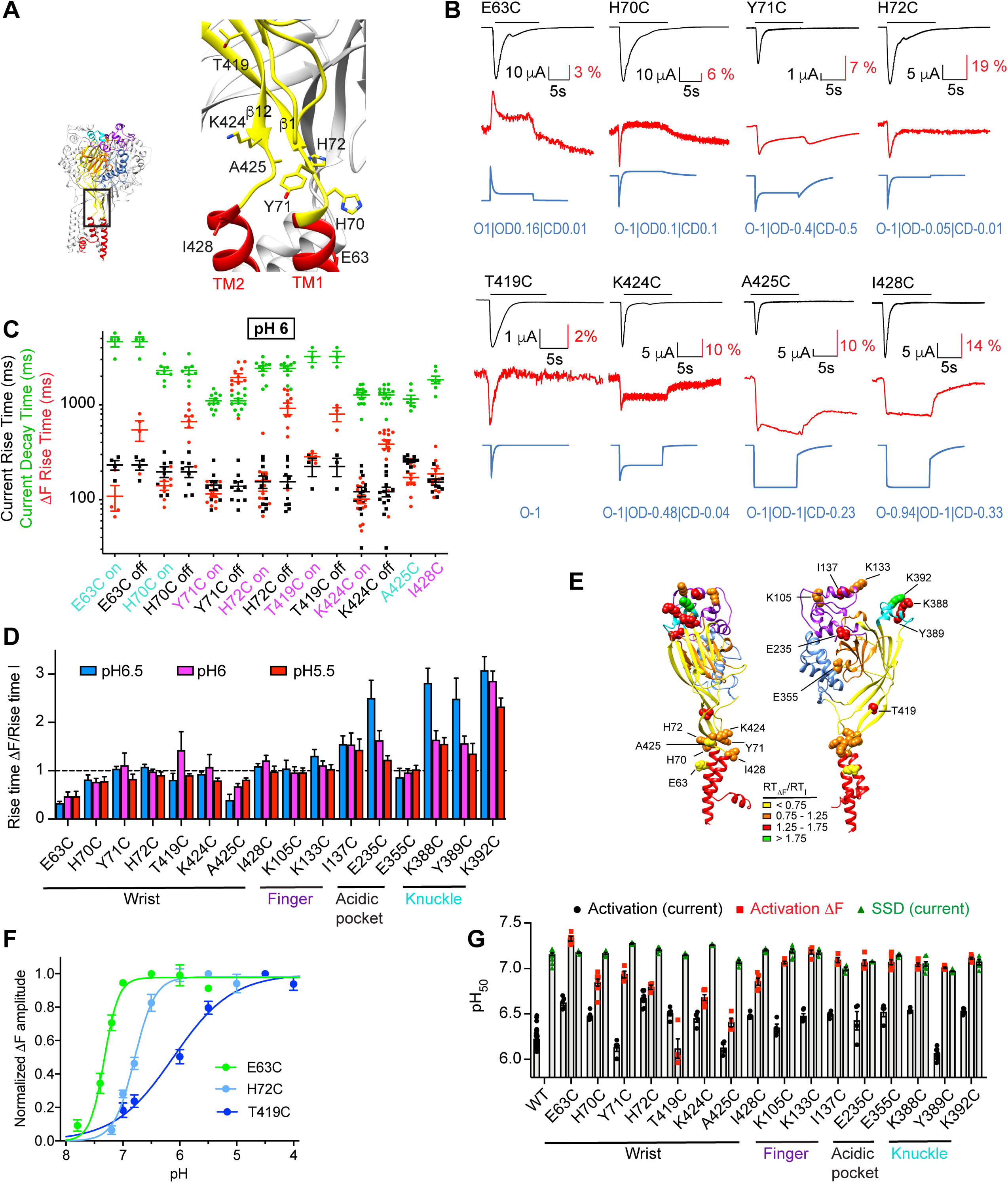
Transient, rapid ΔF signals in the wrist. **(A)** Structural image indicating the positions of the tested mutants. **(B)** Representative current and fluorescence traces of the Cys mutants in the wrist. The conditioning pH between stimulations was pH7.4; the stimulation pH6 was applied as indicated by the horizontal black bars. The blue traces were generated by the kinetic model as described in the legend to Figure 1G. **(C)** The kinetics of current appearance and decay, and of ΔF onset, measured as rise time (RT) or decay time for an activation by pH6.0, are plotted (n=3-14). Since the ΔF signal was transient in many mutants, the kinetics of the ΔF onset (“on”) and decay (“off”) are indicated. The color of the labels of the mutants indicates that the ΔF kinetics are correlated with current appearance (purple) or are faster than current appearance (cyan). **(D)** The RT_ΔF_/RT_I_ ratio is plotted for stimulation pH6.5, 6.0 and 5.5 (n=3-14) of wrist and distant mutants. **(E)** Structural image of ASIC1a (seen from two different sides) indicating the position of the mutants studied in Figures 1 and 2. The color of each residue corresponds to a range of RT_ΔF_/RT_I_ ratio at pH6, as indicated in the figure. **(F)** Experimentally determined pH dependence of the ΔF amplitude for the indicated mutants; n=4-5. **(G)** Summary of pH dependencies of current activation (black), current SSD (green) and ΔF amplitude (red), n = 4-30. In the experiments for ΔF measurements, the conditioning pH was 8.0. Source data are provided in the file Figure2-Source_Data.

To compare the ΔF properties of these mutants with mutations located distantly from the gate, the RT of the ΔF onset was in each experiment normalized to the RT of the current appearance; a ratio < 1 indicates that the ΔF kinetics are faster than the kinetics of current appearance. The RT_ΔF_/RT_I_ ratio, determined at pH6.5, 6.0 and 5.5, was in the range of ∼0.5-1.3 for all wrist mutants (Figure 2D). The RT_ΔF_/RT_I_ ratios are visualized in a color code in the structural image in Figure 2E. The three tested mutants of the finger (K105C, K133C and I137C) and E355C of the acidic pocket displayed also RT_ΔF_/RT_I_ ratios close to 1, while this ratio was higher in the acidic pocket mutant E235C and in the knuckle mutants (K388C, Y389C, K392C) compared to most mutants localized in the wrist at pH6.5 (Two-way ANOVA, Sidak post-test, p<0.05).

The pH dependence of current activation and steady-state desensitization (SSD, corresponding to the closed-desensitized transition), and of the ΔF amplitude were determined (Figure 2F, Figure 2-figure supplement 2), and the pH values of half-maximal effect, pH_50_, are presented in Figure 2G. This shows that even for mutants whose ΔF was associated with channel opening, based on the kinetics and the kinetic model, the pH_50_ of the ΔF signal was in most cases more alkaline than the pH_50_ of current activation, and was for several mutants close to the pH_50_ of SSD. Possible reasons for this shift are discussed below (*Discussion*).

### Fast ΔF signals are associated with channel opening

When the pH is changed from 7.4 to an acidic value, the channel can move from C to either O or CD (Figure 1F). To estimate whether the measured fast ΔF signals might be associated with the C-CD transition (i.e. whether the CD state contributes to this fluorescence), the above-mentioned kinetic model of ASIC1a function (Alijevic et al., 2020) was used to predict the pH dependence of the probability of these different states and of the kinetics of their appearance. Figure 3A plots, based on a step acidification, the normalized peak probability of the four states and of the two gates (O+OD and CD+OD) for the indicated pH range. As expected, the normalized peak probability of the CD state is biphasic (blue in Figure 3A), because of a strong occupancy of the C state at pH ≥ 7, and of the OD state at pH < 6.5. Calculated probability time courses of the different states at different pH conditions further illustrate this biphasic pH dependence of the CD state (Figure 3-figure supplement 1).

**Figure 3.**
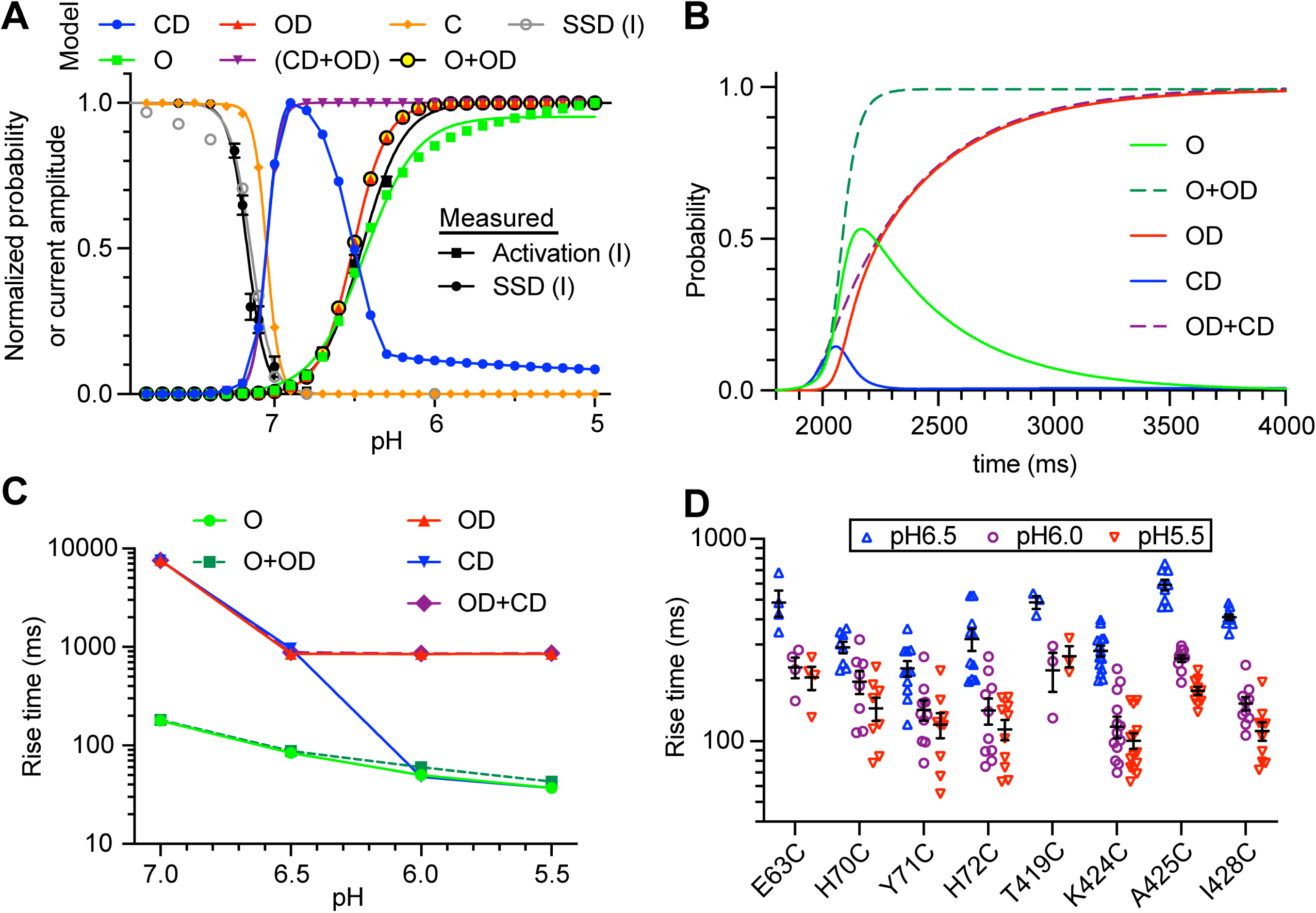
Analysis of pH dependence and kinetics of ΔF signals indicate that rapid ΔF signals are linked to channel opening. (A) Model-generated normalized probability of occupancy of the indicated states (colored traces). The conditions of the simulations were as in Figure 1G, except that the duration of the acidic pulse was 20s. The grey symbols represent the current SSD calculated with the model by applying the same protocol as applied in the experiments (Figure 2-figure supplement 2). The black symbols show the experimental pH dependence of current activation and SSD as determined in Figure 2-figure supplement 2. **(B)** Model-generated time dependence of state occupation upon a pH change from 7.4 to 6.0. Conditioning pH was 7.4; it was changed to pH6.0 at 2s for a duration of 10s. The fall time of the pH change was set to 200 ms. **(C)** Model-generated rise time of the occupancy of the indicated states upon acidification from pH7.4 to the indicated values; model parameters as in B. **(D)** Experimentally determined rise time of the ΔF signal measured for the wrist mutants at the three indicated pH values. Source data are provided in the file Figure3-Source_Data.

The pH dependence of the normalized peak probability of CD, CD+OD and C is shifted alkaline with regard to that of the other states, and is close to the pH dependence of SSD, shown as black and grey symbols for experimental and model data, respectively. The time dependence of the probability of these states for an acidification from pH7.4 to 6.0 (Figure 3B) indicates a rapid increase in the probability of O, O+OD and CD upon acidification. Slow onset kinetics are observed with OD and OD+CD. These kinetics show a certain acceleration with the acidity of the stimulation (Figure 3C) that is also observed with experimental ΔF kinetics (Figure 3D). The model strongly suggests that the fast ΔF signals are associated with opening, since it indicates that a ΔF associated with the CD state would have a biphasic pH dependence, which is clearly not observed in the experiments (Figures 2F-G and 3A). If associated with both the CD and the OD state, the kinetics would be considerably slower than the kinetics of channel opening (Figure 3B). The presented data show so far that rapid conformational changes occur in the wrist and at sites distant from the pore upon acidification. The kinetic ASIC1a model supports the interpretation that these rapid signals are associated with channel opening. To obtain more information on how the rapid conformational changes occurring at the distant sites are transmitted to the gate, we have in the following investigated conformational changes in the palm and nearby domains.

### Slow approaching between the palm β1-β2 linker and the β12 strand

The ΔF signals of A81C, S83C and Q84C, located in the β1-β2 linker, on top of the lower palm β strand 1 (Figure 4A-B), may be due to quenching by Tyr417, which is located on β strand 12. When Tyr417 was mutated to Val (which does not quench the fluorescence) in the background of these Cys mutants, the ΔF signal was strongly decreased in the background of A81C, S83C and Q84C (Figure 4B-C), indicating that ∼70% of the ΔF signal in the single mutants A81C and S83C, and ∼54% in Q84C, was due to a change in distance or orientation relative to Tyr417. The negative ΔF of the single mutants indicates an increased quenching, thus an approaching between the fluorophore attached to the Cys mutants and Tyr417. The ΔF onset kinetics were intermediate between those of current appearance and current decay and in some conditions correlated with current decay (Figure 4D), indicating that the underlying conformational changes may be involved in the induction of ASIC desensitization. The pH_50_ of the ΔF onset was close to that of SSD (Figure 4-figure supplement 1).

**Figure 4.**
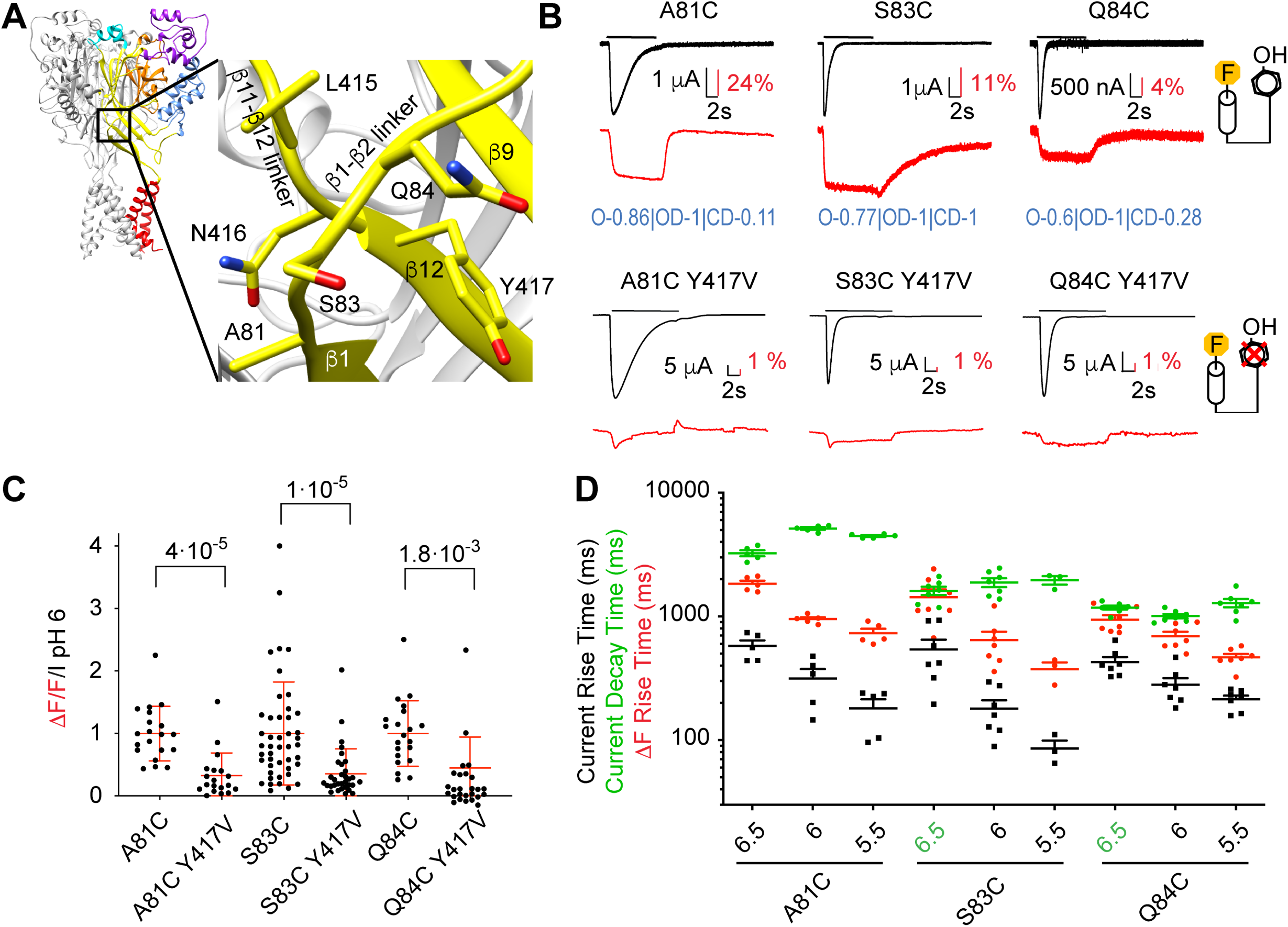
Removal of a fluorescence quenching group strongly reduces β1-β2 linker ΔF amplitude. **(A**) Structural image of ASIC1a (left) with zoom on the lower palm domain (right). **(B)** Representative current and fluorescence traces of single (top) and double (bottom) palm mutants. Conditioning pH7.4 was used, and mutants were stimulated by pH6 for the duration indicated by the horizontal bars. On the right, cartoons of a fluorophore and a quenching Tyr residue illustrate that in the double mutants the quencher was removed. Proportionality factors of the corresponding simulated traces are indicated in blue. **(C)** Normalized ratio of the fluorescence change / total fluorescence, divided by the pH6-induced current amplitude (ΔF/F/I_pH6_) for single and double mutants; Kruskal-Wallis and Dunn’s post-test, n=19-45. Corresponding single and double mutants were measured on the same days, and for each experimental day, the ΔF/F/I_pH6_ ratio of each cell of a given single/double mutant pair was divided by the average ΔF/F/I_pH6_ ratio of the single mutant of that day. **(D)** Current RT and decay time and ΔF RT of single mutants obtained at pH6.5, 6.0 or 5.5 (n=3-7). A green pH indication indicates that the ΔF onset kinetics are correlated with current decay. Source data are provided in the file Figure4-Source_Data.

### Conformational changes in the β1-β2 linker and the β5-β6 loop precede desensitization

These double mutants were then used as a basis for placing Trp, a strong fluorescence quencher, at different positions in the proximity of the fluorophore docking positions A81C, S83C or Q84C, to monitor additional distance changes. Studies with different fluorophores indicated quenching by Trp at distances ≤ 15 Å (Mansoor et al., 2002, Pantazis et al., 2018). The β5-β6 linker of the β-ball runs almost parallel to the membrane surface, above the β1-β2 linker. In the direction from Pro205 to Met210 it approaches a subunit interface (Figure 5A) (Lynagh et al., 2018, Bignucolo et al., 2020). Nine mutants were identified in which the placement of a Trp in the β5-β6 palm loop generated fluorescence signals (Figure 5A-B). Most triple mutants showed normal transient currents. Only those containing mutations of K208, T209 and M210, and L207 in the background of A81C/Y417V had a slowed or disrupted desensitization (Figure 5B-C). The ΔF signals at pH6.0 were in most cases sustained and of a single component, except for the ΔF signals of Q84C/Y417V/P205W and A81C/Y417V/L207W, which appear to be the sum of an early negative and slightly delayed positive ΔF component (Figure 5B). A control stimulation protocol in which the channels are exposed to pH6.7 (which desensitizes the channels) before stimulation by pH6 was used to test whether a signal may potentially be non-specific (*Materials and Methods*, Supplementary Table S3). ΔF components identified as potentially non-specific are marked “ns” in Figure 5B and were not further considered for the analysis. The ΔF kinetics of most mutants were slower than those of current appearance (Figures 5C, 5-figure supplement 1), indicating that the underlying conformational changes are likely preparing or associated with desensitization. The ΔF traces of most mutants were well reproduced by the kinetic model, except for Q84C/Y417V/P205W and A81C/Y417V/L207W (Figure 5-figure supplement 2). Most mutants had highest proportionality factors with the OD, and smaller with the O state. The VCF analysis of these fluorophore-quencher pairs indicated thus 9 distance changes, 2 of them associated with opening, 7 preparing desensitization (Table 2). These findings were then used to obtain information on conformational changes, as shown further below.

**Figure 5.**
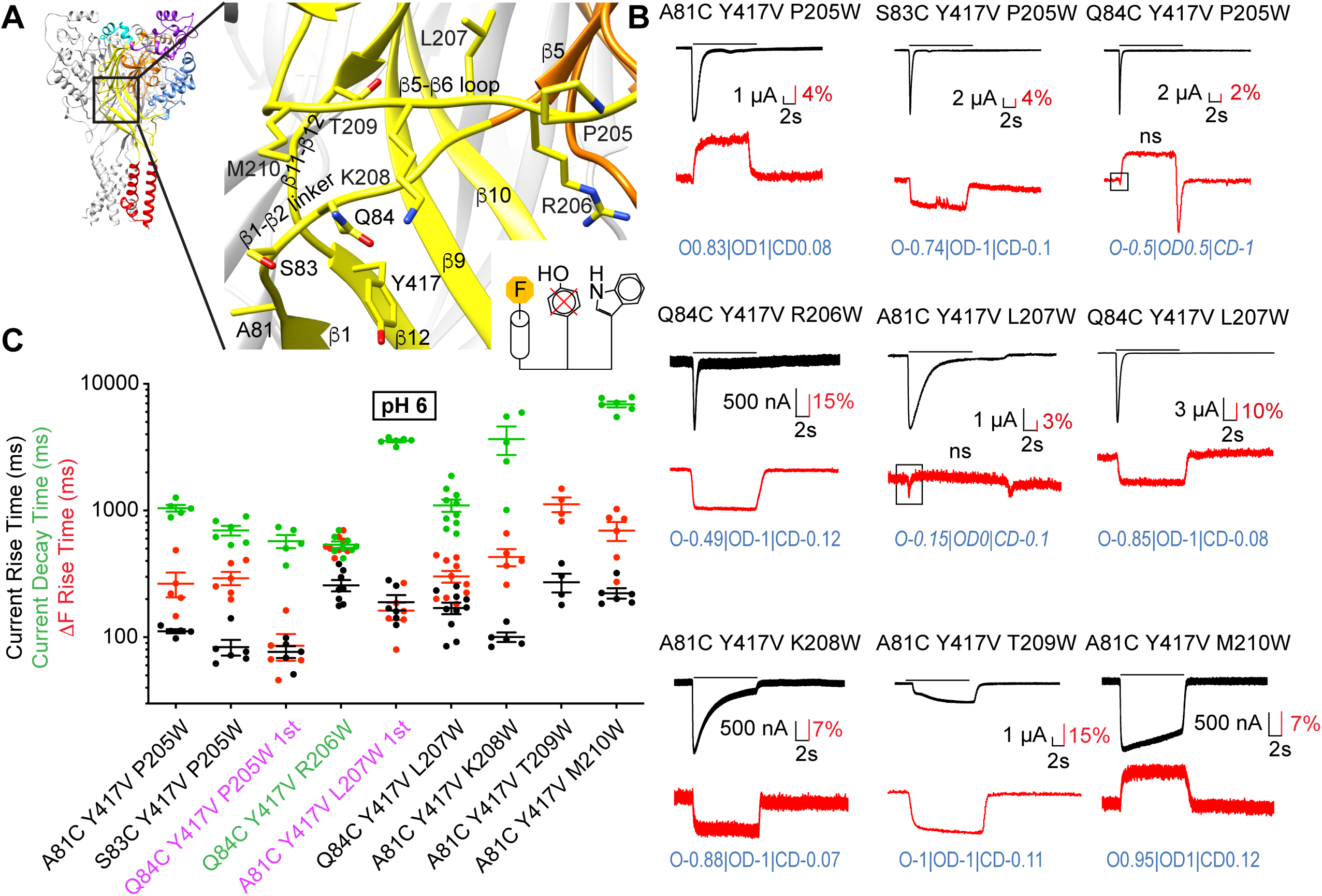
Slow structural rearrangements in the palm domain. **(A**) Structural image of ASIC1a (left) with a zoom on the palm (right), showing the position of mutations studied in this figure. Bottom, cartoon illustrating the approach used for these triple mutants, with the fluorophore (“F”, on the left), the original quenching group Try417 (center) mutated, and a new quenching Trp (right) introduced at a different position. **(B)** Representative current and fluorescence traces of the triple mutants. Conditioning pH7.4 was used, and mutants were stimulated by pH6 for the duration indicated by the horizontal bars. The black frames in some ΔF traces highlight the first ΔF component. “ns” indicates that this ΔF component was potentially non-specific (Table S3). Proportionality factors of the corresponding simulated traces are indicated in blue. **(C)** Current RT and decay time and ΔF RT obtained at pH6, n=4-10. The color of the labels of the mutants indicates that the ΔF onset kinetics are correlated with current appearance (purple) or decay (green; *Materials and Methods*). Source data are provided in the file Figure5-Source_Data.

**Table 2.** Summary of VCF and SMD analyses.

| Residue1 | Residue2 | VCF signal |  |  | SMD simulations |  |  |  |  |  | Validation |
| --- | --- | --- | --- | --- | --- | --- | --- | --- | --- | --- | --- |
|  |  | Comp. | Signal | Ass. | Deviation<br>(Å) |  |  | Force<br>(kJ mol<sup>-1</sup> Å<sup>-1</sup>) |  |  |  |
| Closed-to-open |  |  |  |  |  |  |  |  |  |  |  |
| Q84 | P205 | 1<sup>st</sup> | neg | o |  |  |  |  |  |  |  |
| A81 | L207 | 1<sup>st</sup> | neg | o | 2.4 | ± | 1.3 | -7.1 | ± | 3.8 | no |
| S83 | T289 |  | **neg** | o | 1.3 | ± | 0.5 | -3.8 | ± | 1.4 | yes |
| S83 | Q358 | 1<sup>st</sup> | **pos** | o | -0.6 | ± | 0.4 | 1.9 | ± | 1.1 | yes |
| S83 | E359 |  | *neg* | o | 1.2 | ± | 0.3 | -3.5 | ± | 0.9 | yes |
| A81 | L369 | 1<sup>st</sup> | **neg** | < o | 2.8 | ± | 0.5 | -8.4 | ± | 1.5 | no |
| Open-to-desensitized |  |  |  |  |  |  |  |  |  |  |  |
| A81 | Y417 |  | **neg** | i | 2.2 | ± | 0.8 | -6.5 | ± | 2.3 | no |
| S83 | Y417 |  | **neg** | i | 1.1 | ± | 0.3 | -3.5 | ± | 1.3 | yes |
| Q84 | Y417 |  | neg | i | 1.4 | ± | 0.9 | -4.1 | ± | 2.6 | yes |
| A81 | P205 |  | pos | i |  |  |  |  |  |  |  |
| S83 | P205 |  | neg | i | -0.1 | ± | 0.5 | 0.3 | ± | 1.4 | yes |
| Q84 | R206 |  | neg | d | 1.2 | ± | 1.3 | -3.5 | ± | 4.0 | yes |
| Q84 | L207 |  | neg | i | 2.0 | ± | 1.4 | -6.1 | ± | 4.1 | no |
| A81 | K208 |  | **neg** | i | 1.9 | ± | 0.4 | -5.8 | ± | 1.2 | no |
| A81 | T209 |  | **neg** | i |  |  |  |  |  |  |  |
| A81 | M210 |  | *pos* | i | -2.6 | ± | 0.7 | 7.8 | ± | 2.2 | no |
| A81 | T289 |  | *neg* | i | 1.4 | ± | 0.3 | -4.3 | ± | 0.9 | yes |
| Q84 | T289 |  | **neg** | i | 1.9 | ± | 0.4 | -5.8 | ± | 1.3 | no |
| S83 | D357 | 1<sup>st</sup> | **neg** | d | 0.6 | ± | 0.7 | -1.7 | ± | 2.2 | yes |
| S83 | Q358 | 2<sup>nd</sup> | neg | d | 1.8 | ± | 0.5 | -5.3 | ± | 1.5 | no |
| A81 | L369 | 2<sup>nd</sup> | *pos* | d | -0.1 | ± | 0.5 | 1.4 | ± | 1.4 | yes |
| S83 | L369 | 2<sup>nd</sup> | pos | i | -2.2 | ± | 0.5 | 6.7 | ± | 1.6 | no |

### Detection of fast conformational changes in palm-thumb loops

To follow the kinetics of distance changes to close residues of a neighboring subunit, fluorophore docking at A81C, S83C or Q84C, in the background of the Y417V mutation, was combined with positioning of the quenching group in three different sub-domains, at different distances from the plasma membrane (Figure 6A): 1) close to the β-turn (‘T289W; the prefix’ indicates a residue of a neighboring subunit), 2) in the palm β10 strand (’L369W), and 3) at the end of the thumb helix α5 and in the linker between the thumb α5 helix and the palm β strand 10 (‘D357W, ‘Q358W, ‘E359W; “α5-β10 loop”). The distances compatible with quenching in these triple mutants are those between subunits (Figure 6A). These triple mutants produced normal transient pH-activated ASIC currents, with the exception of S83C/Y417V/D357W, which showed only a very small current with a transient and a sustained component (Figure 6B). The ΔF kinetics of four out of eight of these mutants were correlated with the kinetics of current appearance or were faster, indicating that at least one of the partners (S83/’T289, S83/’Q358, S83/’E359, A81/’L369) moves with the speed of activation (Figures 6B-C and 6-figure supplement 1). Most ΔF traces could be well fitted with the kinetic model (Figure 6-figure supplement 2) indicating generally high proportionality factors for OD and O. This set of VCF measurements provided information on 4 distance changes occurring in the C-O transition, and 6 in the O-D transition.

**Figure 6.**
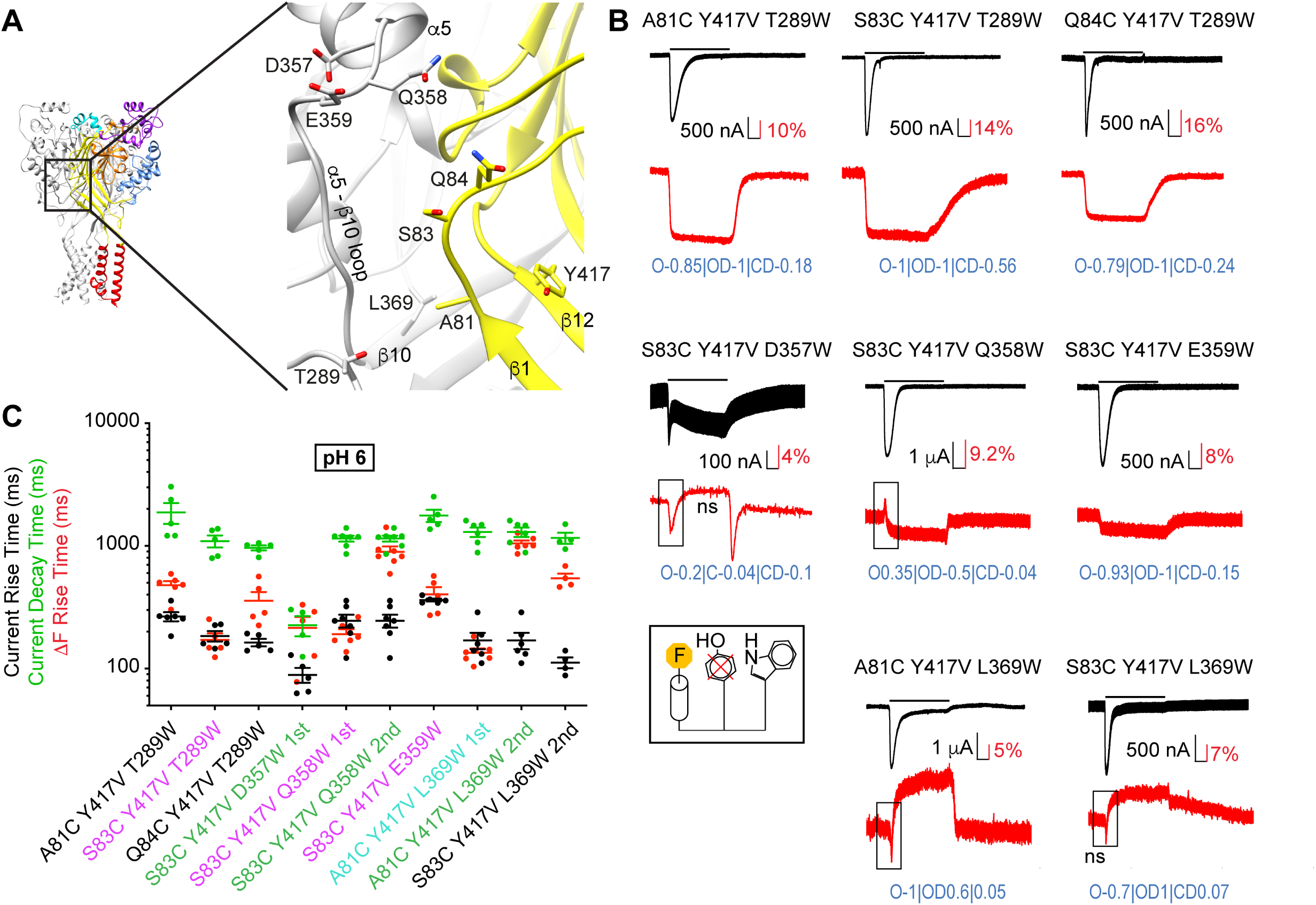
Fast structural rearrangements in the palm-thumb loops. **(A**) Structural image of ASIC1a (left) with a zoom on the palm (right), showing the position of mutations studied in this figure. **(B)** Representative current and fluorescence traces of the triple mutants. Conditioning pH7.4 was used, and mutants were stimulated by pH6 for the duration indicated by the horizontal bars. The black frames in some ΔF traces highlight the first ΔF component. “ns” indicates that this ΔF component was potentially non-specific (Table S3). Proportionality factors of the corresponding simulated traces are indicated in blue. **(C)** Current RT and decay time and ΔF RT obtained at pH6, n=4-7. The color of the labels of the mutants indicates that the ΔF onset kinetics are correlated with current appearance (purple) or decay (green) or are faster than current appearance (cyan; *Materials and Methods*). Source data are provided in the file Figure6-Source_Data.

Analysis of the pH dependence showed that the ΔF was more pH-sensitive than was the current activation in the majority of the palm triple mutants (Figure 6-figure supplement 3). There was no trend of mutants with faster ΔF kinetics having ΔF pH_50_ values closer to their pH_50_ of current activation (Figure 6-figure supplement 4).

### Structural interpretation and validation of VCF-predicted distance changes by Steered Molecular Dynamics simulations

The VCF measurements of fluorophore-quencher pairs indicate whether the residues of a given pair move towards or away from each other during channel opening or desensitization. They do not indicate whether the different predictions are equally reliable, and whether they are compatible with the structures. Steered molecular dynamics (SMD) simulations were therefore used to determine whether predicted distance changes were likely to occur, and to provide structural interpretation of the VCF-based predictions. The SMD simulations were carried out starting either from a structural model representing the closed state (for ΔF signals associated with the C-O transition) or the open state (O-D transition), embedded in a 1-palmitoyl-2-oleoyl-sn-glycero-3-phosphocholine (POPC) bilayer. In a MD simulation, the atom positions are calculated over time by applying the classical equations of motion on the ensemble of particles exposed to a potential, which integrates the known features of interactions into an ensemble of equations and parameters called a force field. In a SMD simulation, the force field is extended by additional pulling forces applied between pairs of atoms or between the centers of mass of atom groups. A combination of harmonic potential and a target distance was applied (see *Materials and Methods)* to either bring pairs of residues closer to each other or move them away from each other, according to the VCF predictions. The target distance was set to the distance in the structural model plus or minus 5Å for residues predicted to move away from or towards each other, respectively. The magnitude of the pulling force depended on its effectiveness: as long as the target distance was far from reach, the magnitude of the force remained high.

### Complex intersubunit distance changes are associated with channel opening

The VCF analysis indicated two intrasubunit and four intersubunit distance changes occurring before or with channel opening (Figure 7-figure supplement 1). Of these, the pair Gln84-Pro205 had only a small ΔF amplitude and was therefore not further considered for structural interpretation, leaving five constraints for the SMD simulation starting from the closed state. Figure 7A presents in the upper panel the distance between the Ser83 and ‘Q358 side chains as a function of time during the SMD simulation, while the lower panel shows the pull force for the same pair. In this pair, the target distance (11.5 Å, horizontal blue line in Figure 7A) was approached very rapidly, and the pulling force decreased with a similar time course. For the pair Ala81-Leu207 (Figure 7B) the target distance was only partially approached during the simulation, and the pull force remained larger than 7 kJ/mol/Å. The MD trajectories were considered as having reached the target distance of a given pair, and therefore the VCF-predicted distance change was validated, if the distance reached during the last 40ns of the SMD simulation was within 2.2 Å of the expected target and if the remaining applied force was smaller, in absolute value, than 6 kJ/mol/Å. Table 2 reports for each pair the deviation from the expected target, the pull force exerted during the same time interval, and whether the SMD validated the VCF-predicted distance change.

**Figure 7.**
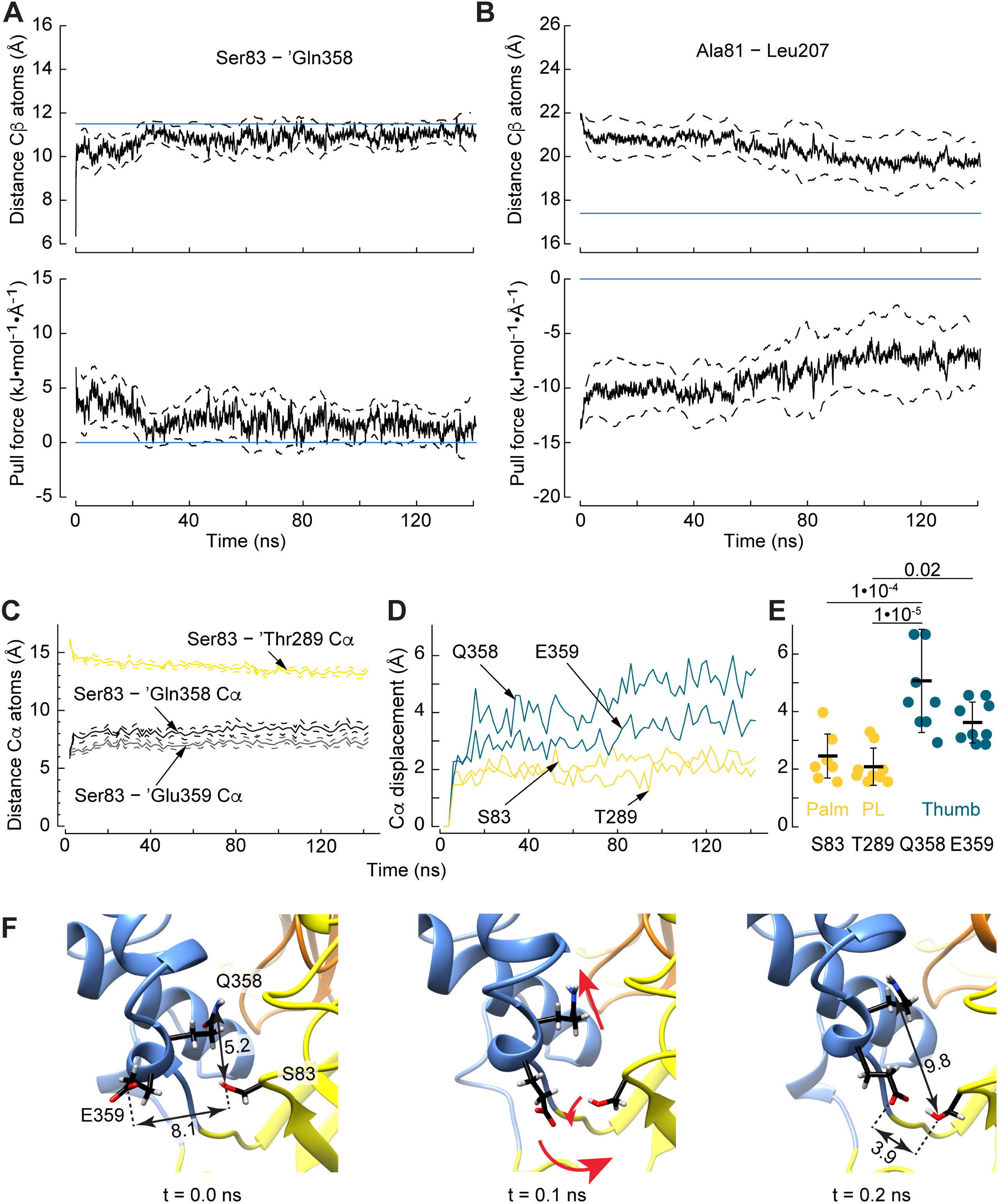
Steered Molecular Dynamics (SMD) simulations identify the most reliable VCF-predicted conformational changes and describe movements during the channel opening. SMD trajectories started from the closed structural model, showing the pair Ser83-Gln358 **(A)** in which the distance goal (upper panel) was reached, and the pull force (lower panel) vanished, and the pair Ala81-Leu207 **(B)** in which the difference between the target and the measured distance between the two residues, and the pull force remained high. The blue horizontal lines indicate the target distances (upper panels) and the absence of pull force (lower panels). **(C)** Time series of distances between Cα pairs of atoms. The initial distances between Ser83 and ‘Thr289, ‘Gln358 and ‘Glu359 were ≍ 16, 6 and 6 Å, and the distances at the end of the trajectories were ≍ 13, 9 and 7 Å. Means and SEM are shown, n = 9. **(D)** Time series of the Cα displacement of the indicated residues, colored according to their location in the palm, the palm-thumb linker (PL) or the thumb. For clarity, only the means from the n = 9 trajectories are shown. **(E)** Means and SD of the Cα displacements. The first 100 ns of simulations were discarded for this analysis (One-way ANOVA and Tukey Honest Significant Differences post-test, n = 9). **(F)** The simultaneous approaching of Ser83 to ‘Glu359 and distancing between Ser83 and ‘Gln358 during the closed to open transition is possible due to rotation of the ‘Glu359 sidechain and other local movements (red arrows). These reorientations and translations result in the switch of the Ser83-interacting residue from ‘Gln358 to ‘Glu359, as shown in three molecular representations of the protein conformation at successive times of a SMD trajectory. The distances between the OG atom of Ser83 and the CD atoms of Glu359 and Gln358 are indicated. Source data are provided in the file Figure7-Source_Data.

Following these criteria, three out of the five tested movements were validated, an approaching between Ser83 and 1) ‘Thr289 and 2) ‘Glu359, and 3) a distance increase between Ser83 and ‘Gln358 (Table 2, Figure 9B). VCF experiments do not indicate which of the two residues within a pair moves the most, and whether only the side chain, or also the backbone moves. Of the three pairs, the Cα distance increased and decreased, respectively, for Ser83-’Gln358 and Ser83-’Thr289 (Figure 7C), supporting the interpretation that in these pairs the distance between the backbone changed, while in contrast the Cα distance of the Ser83-’Glu359 did not change. To detect movements of the backbone, the average displacement of the Cα atoms during the SMD was extracted. The SMD attributed significantly higher Cα movements to the thumb residues Gln358 and Glu359 than to Ser83 and ‘Thr289 (Figures 7D-E). This indicates that in the S83-’Thr289 pair the Cα atoms move towards each other, with a similar, small displacement, while in the 83-’Gln358 and 83-’Glu359 pairs mostly the thumb residues move. The simultaneous increase in the Ser83-‘Gln358, and decrease in the Ser83-‘Glu359 distance may appear incompatible at first view. However, inspection of the SMD trajectory showed that the side chain of ‘Glu359 is oriented away from the β1-β2 linker and Ser83 is located within electrostatic interactions of ‘Gln358 in the initial structure (Figure 7F). The applied pulling forces during the SMD resulted in a fast rotation of Glu359 towards Ser83, whereas ‘Gln358 was pushed away. In a recent study we observed that the distance between ‘Glu359 and the neighboring palm domain was correlated with the pH dependence of activation (Bignucolo et al., 2020), suggesting that facilitating the decrease of this distance would enhance the activation probability.

**Figure 9.**
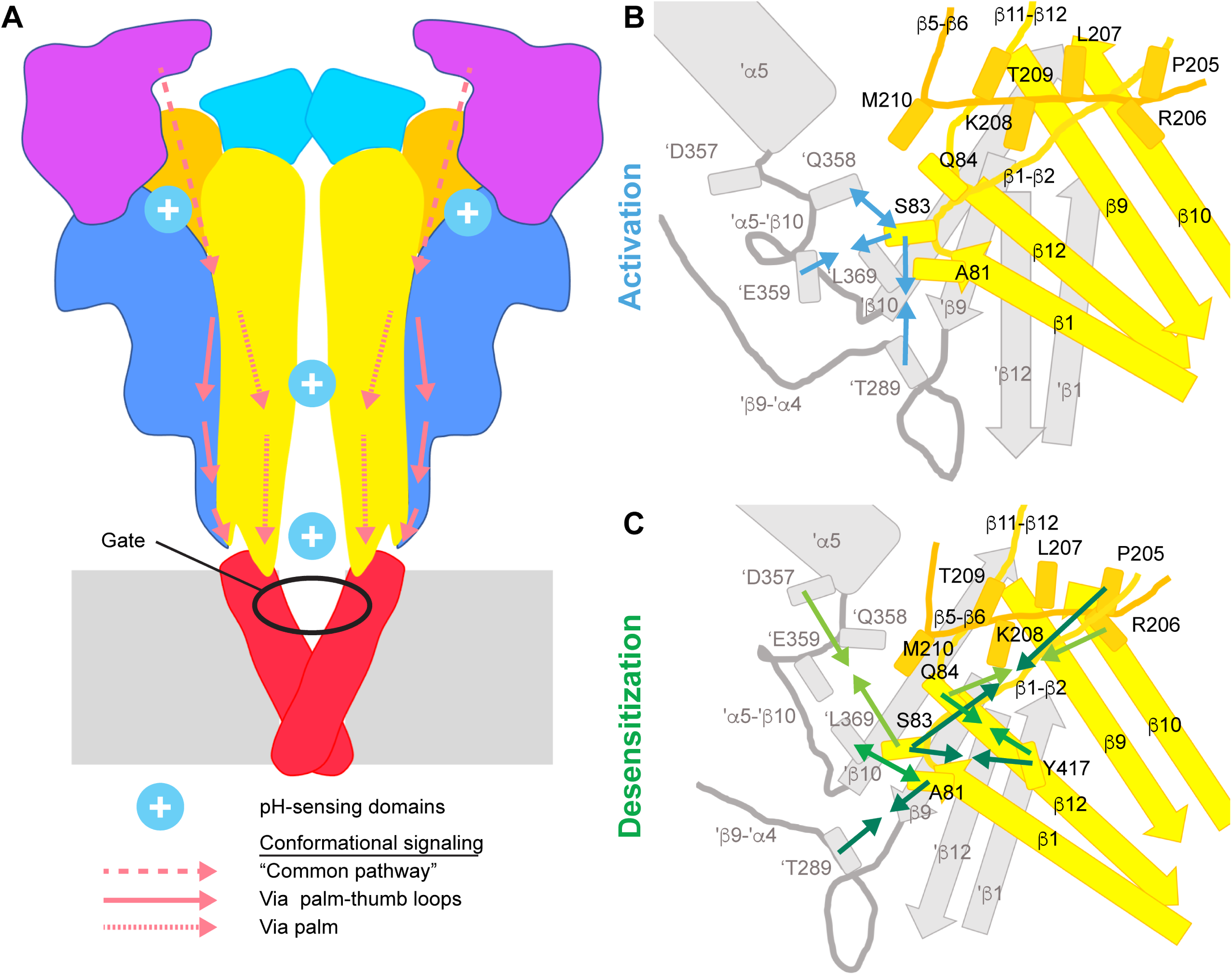
Conformational changes leading to activation and desensitization. (**A**) Activation signaling pathways in ASIC1a. Cartoon of the ASIC structure, in which the predicted pH-sensing regions are highlighted (only two of the three acidic pockets are indicated) and hypothesized signaling pathways for channel activation from distal regions to the gate are indicated; dashed arrows, common pathway, dotted arrows; palm pathway; solid arrows, palm-thumb loop pathway, with the β-turn interacting with the upper end of the TM1. (**B**) and (**C**), Cartoons indicating the conformational changes in the palm and palm-thumb loops predicted by VCF that were validated by SMD simulations to occur during ASIC activation (**B**) and desensitization (**C**). The lower palm domain of one subunit is shown in yellow, structural elements of a neighboring subunit are shown in grey. Arrows that point toward each other indicate an approaching between residues, while arrows pointing away from each other indicate an increase in distance.

### An approaching between subunits associated with desensitization

The VCF results predicted 16 specific distance changes between residue pairs with ΔF kinetics that were either intermediate between those of opening and desensitization or correlated with desensitization (Figure 8-figure supplement1). Of these, two pairs were not considered for the analysis of distance changes, Ala81-Pro205 because of the high distance (∼24Å), and Ala81-Thr209 because the A81C/Y417V/T209W current did not desensitize. In the SMD simulations starting from the open state conformation, constraints between 14 residue pairs were therefore included. This analysis, using as thresholds 2.2 Å and 5 kJ/mol/Å (*Materials and Methods)*, validated four intrasubunit and three intersubunit distance changes, indicating an approaching between β1-β2 linker residues and 1) the neighboring thumb and ‘β9-’α4 loop, 2) the β12 strand and 3) the distal β5-β6 linker, and a distance increase between Ala81 and ‘L369 of the neighboring palm (Table 2, Figure 9C).

Of these VCF-predicted and SMD-validated distance changes, the following appeared to contradict the predictions that are based on the published structures. The approaching between ‘Thr289 and the neighboring β1-β2 linker, already observed during opening, was continued during desensitization, whereas a comparison of the crystal structures suggests that they move away from each other. Also, the validated distance increase between Ala81 and ‘Leu369 of the neighboring palm contradicts the structure predictions. For these two pairs, the movement of the Cα atoms goes in the same direction as that of the side chains (Figure 8A), supporting thus a movement of the backbone. The displacement analysis suggests that Ala81 as well as ‘Thr289 both move substantially, contrary to Leu369 (Figure 8B-C). Thr289 is located close to the β-turn in the β9-β4 loop. In cASIC1a, Thr289 is replaced by an Asp residue, which could engage into different interactions with the neighboring Lys292 (Lys291 in hASIC1a). In addition, this loop is relatively poorly resolved in the crystal structure, since residues 297 and 298 (hASIC1a-295 and −296) are missing, thus possibly decreasing the reliability of some structural distances. Our analysis does not reveal any obvious explanation for the contradiction related to the Ala81 - ‘Leu369 pair. Leu369 is located within a narrow pocket of hydrophobic residues, in line with its low mobility in the simulations (Figure 8B-C). Since Ala81 is located close to Leu415 and Asn416 that undergo an isomerization during desensitization (Baconguis and Gouaux, 2012, Rook et al., 2020), it may be involved in conformational changes linked to this isomerization.

**Figure 8.**
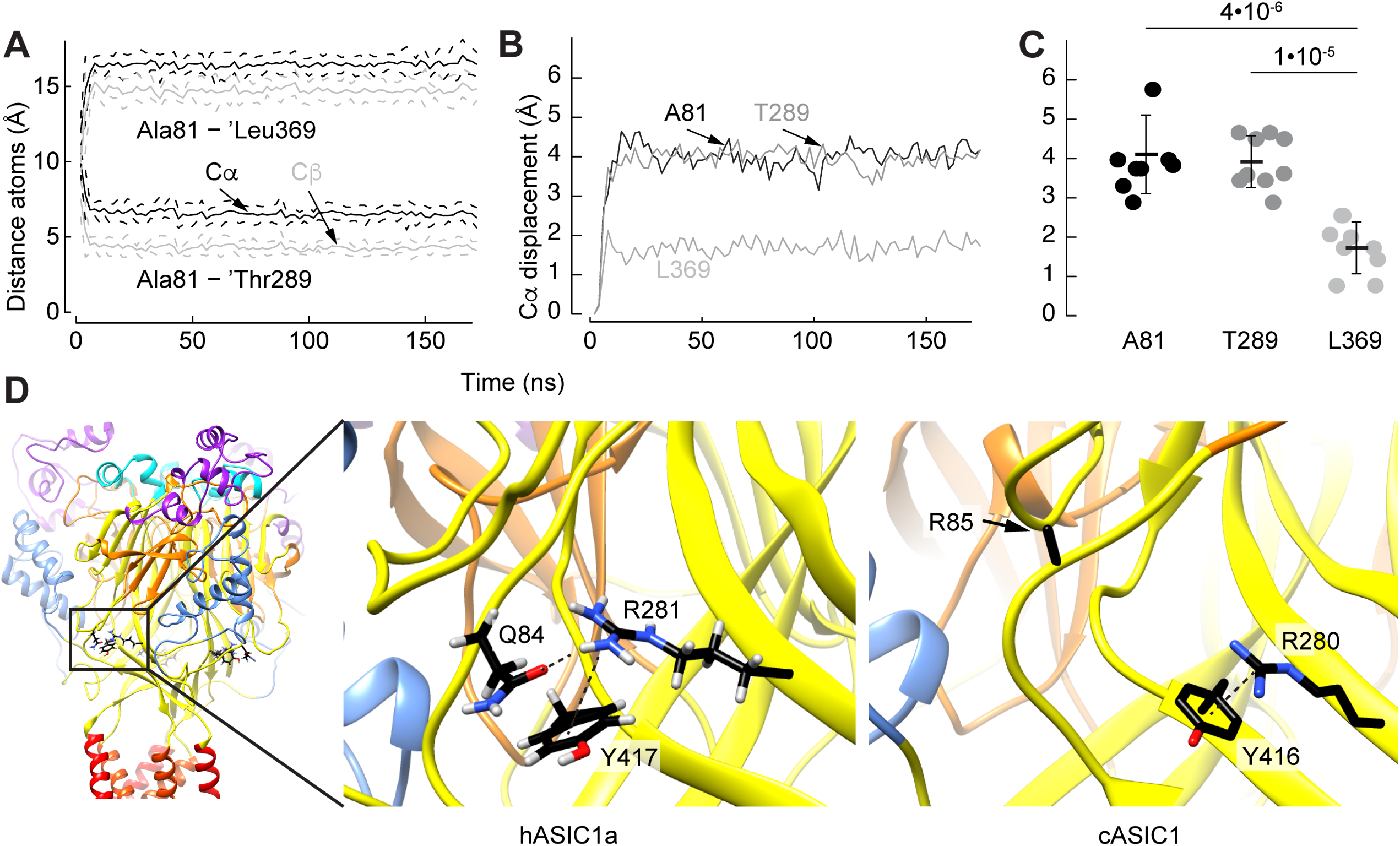
SMD simulations describe movements during the channel desensitization and explain apparent contradictions. **(A)** Time series of distances between Cα (black) and Cβ (grey) pairs of atoms. The initial distances between the Cα atoms of Ala81and ‘Thr289 and ‘Leu369 were ≍ 10 and 12 Å, and the distances at the end of the trajectories were ≍ 6 and 16 Å, respecively. Means and SEM are shown, n = 9. **(B)** Time series of the Cα displacement of Ala81, Thr289 and Leu369, mean values, n=9. **(C)** Means and SD of the Cα displacements, starting at t = 100 ns. (One-way ANOVA and Tukey Honest Significant Differences post-test, n = 9). **(D)** The hASIC1a, but not the cASIC1a structure, favors the approaching between Gln84 and Tyr417 during the open to desensitized transition. The homologous residues in cASIC1a are Arg85 and Tyr416. Left, structural image of hASIC1a. The frame indicates the magnified area shown in the center panel. Center, molecular representation from a SMD trajectory of hASIC1a showing that Tyr417 forms a cation-π interaction with Arg281, which also undergoes a hydrogen bond with Gln84 during the simulation. Right, In the cASIC structure (PDB entry 4NTW) of the corresponding area, Tyr416 also forms a cation-π interaction with Arg280, which is homologous to hASIC1a-Arg281. However, in this case, Arg85 is not likely to approach this positively charged area. Note that the side chain of Arg85, except Cβ, is absent from this open as well as from the desensitized structure (PDB code 4NYK). Electrostatic interactions are symbolized with black dashed lines. Source data are provided in the file Figure8-Source_Data.

The open and desensitized structural models show no difference in distance between the β1-β2 linker and the distal residues Pro205 and Arg206 of the β5-β6 linker, nor between Gln84 and Tyr417 (Table 1), while validated VCF predicts in these cases an approaching. These differences arise likely from the fact that the residue homologous to hASIC1a-Gln84 is an Arg in cASIC1a (Arg85). In cASIC1a, the approaching between the β1-β2 and the distal β5-β6 linker would bring Arg85 and Arg207 (homologous to hASIC1a-Arg206) close to each other, which is unlikely to occur due to the repulsion between Arg residues (Figure 8D). In the cASIC1a open structure, Arg280 (Arg281 in hASIC1a) forms a cation-π interaction with Tyr416, which is homologous to Y417 in hASIC1a. Stabilized in its orientation by this interaction, it points towards the putative side chain of Arg85, thus preventing a favorable approaching between Arg85 and Tyr416. This barrier is absent in the hASIC1a structure, since the carboxyamide group of Gln84 can rotate and form a hydrogen bond between its oxygen atom and Arg281, thus favoring the approaching between Gln84 and tyr417.

**Table 1.** Distances between Cys / quencher pairs.

| Residue1 | Residue2 | Distance (from structure) |  |  |
| --- | --- | --- | --- | --- |
|  |  | Average | Difference |  |
|  |  |  | open-closed (Å) | desensitized-open (Å) |
| A81 | Y417 | 8.4 | 0.3 | -1.5 |
| S83 | Y417 | 10.9 | 0.0 | -2.7 |
| Q84 | Y417 | 10.5 | 0.0 | -0.4 |
| A81 | P205 | 23.5 | 0.1 | -0.5 |
| S83 | P205 | 17.4 | -0.4 | -0.4 |
| Q84 | P205 | 13.0 | -0.1 | -0.6 |
| Q84 | R206 | 11.0 | -0.2 | 0.0 |
| A81 | L207 | 19.6 | 0.4 | -2.1 |
| Q84 | L207 | 10.6 | -0.2 | 0.7 |
| A81 | K208 | 15.0 | 0.3 | -1.3 |
| A81 | T209 | 13.1 | -0.2 | -3.1 |
| A81 | M210 | 11.7 | -0.4 | -3.0 |
| A81 | T289 | 8.5 | 0.2 | 2.2 |
| S83 | T289 | 12.4 | -2.0 | -0.9 |
| Q84 | T289 | 16.2 | -0.8 | -1.7 |
| S83 | D357 | 11.5 | 3.6 | -1.7 |
| S83 | Q358 | 7.1 | 4.5 | -0.5 |
| S83 | E359 | 8.8 | 3.0 | -1.0 |
| A81 | L369 | 10.9 | -2.4 | -2.1 |
| S83 | L369 | 18.2 | -3.7 | -0.9 |

The distances were calculated between the $\beta$ -carbon residues of the indicated residues from structural models of the closed ((Yoder et al., 2018) PDB code 5WKU), Mit-toxin-opened ((Baconguis et al., 2014) 4NTW) and desensitized state ((Gonzales et al., 2009) 4NYK) of hASIC1a in Chimera (Pettersen et al., 2004). The distances were measured in all three subunits, and the average was calculated. "Average distance" is the average of the distance measured in the structural models corresponding to the three functional states. The differences between these distances are presented in a way that negative values indicate a shorter distance in open relative to the closed, or desensitized relative to the open conformation.

## Discussion

We show here that conformational changes related to channel opening occur at the same time in the extracellular pore entry and in more distant sub-domains, such as the finger, acidic pocket, palm and thumb-palm loops, supporting the view that pH sensing at peripheral sites contributes to ASIC activation. Our combined VCF and SMD analysis indicates that fast conformational changes tend to occur more in the two thumb-palm loops, while slower changes occur in the palm. The lower thumb-palm loop contains the β-turn that interacts with the TM1 and may, via this interaction, lead to the opening of the channel gate (Figure 9A).

### Association of ΔF signals with functional transitions

To test for an association between ΔF signals and functional transitions, the kinetics of current and ΔF signals were compared. This showed, comparable to many other VCF studies, that some ΔF signals are timely associated with defined functional transitions, while many mutants showed intermediate kinetics, likely because they report conformational changes that are not directly associated with the channel gate but may be part of processes that precede and lead to the functional transition. In addition to this analysis, we have applied kinetic modeling that assumed that the fluorescence signal is a linear combination of the probabilities of finding the ASICs in individual functional states. Although this simple model does not consider any intermediate states and. is based on WT ASIC1a, it was able to reproduce most of the measured ΔF signals rather well. The scaling factors in these models suggested for most mutants that the open and open-desensitized states were the principal contributors to the ΔF signal.

For the interpretation of VCF data, potential limitations have to be taken into consideration. VCF is carried out with mutants, and although mutants are selected that have similar properties as the WT, there are differences with regard to the kinetics and the ligand concentration dependence. In this study, ΔF kinetics were directly compared to the current kinetics, therefore the conclusions were not influenced by kinetic differences of the mutants to the WT. VCF measurements were done in the pH range 6.5-5.5 and were qualitatively similar over this range. Since for the vast majority of the tested mutants the pH_50_ shift relative to the WT was <0.5 pH units, these shifts do not influence the conclusions.

Experiments with ultrarapid perfusion systems have shown that ASICs activate with time constants of the order of 10 ms (Bassler et al., 2001, Sutherland et al., 2001, Alijevic et al., 2020). In the measuring chamber used in this study, the RT of solution change was ∼300ms (*Materials and Methods)*. The measured kinetics were therefore limited by the speed of solution change, and since the measured RTs of many mutants were faster than 300ms, peak amplitudes were probably reached before the solution was completely exchanged (*Materials and Methods)*. In each experiment and for each condition, the kinetics of ΔF and current were directly compared, providing thus correct conclusions of the ΔF kinetics relative to the current kinetics, and therefore a valid attribution of the ΔF signals to functional transitions. In the interpretation of the data, it has to be considered that the absolute speed of the measured kinetics was limited by the speed of the solution change, and that the actual pH of the measurement of the kinetics was slightly more alkaline than that of the test solution.

### Dissociation of ΔF and current pH dependence

The pH dependence of the ΔF signal was in most mutants, even mutants associated with activation, shifted to more alkaline values than the pH dependence of current activation, and closer to the pH dependence of SSD. Such a shift has been observed in previous studies (Vullo et al., 2017, Bonifacio et al., 2014). It was observed here even with the non-desensitizing mutant A81C/Y417V/T209W, thus it does not indicate a link to desensitization. It is thought that in ASICs, as in other ion channels, conformational changes in different parts of the channel will eventually lead to the opening of the channel pore. The pH dependence of the current reflects the pH dependence of the ultimate steps in this pathway. It is possible that conformational changes that do not belong to these ultimate steps have a different pH dependence. The pH_50_ values depend on the two ends of the scale, the most alkaline pH at which a ΔF signal or a current is detected, and the pH at which its increase saturates. ΔF signals appear to saturate in many mutants at less acidic pH than the current. The observed alkaline shift in the pH dependence of fluorescence is reminiscent of the hyperpolarization shift of the fluorescence relative to the current voltage dependence of K^+^ channels (Mannuzzu et al., 1996, Cha and Bezanilla, 1997). A divergence of the concentration dependence between the ΔF signals and the current was also observed for other ligand-gated channels for fluorescence signals associated with activation (Dahan et al., 2004, Pless and Lynch, 2009).

### Structure-derived rearrangements in the palm and wrist

Structure comparisons indicate that ASIC opening is associated with a collapse of the central vestibule - enclosed by the upper parts of the lower palm β sheets - and at the same time an expansion of the lowest end of the lower palm, together with an iris-like opening of the gate located in the transmembrane domains (Yoder et al., 2018). These rearrangements are accompanied by a displacement of the β-turns. The collapse of the central vestibule continues during the open-desensitized transition. It was proposed that during desensitization the pore would be uncoupled from this continued movement, allowing the gate to relax back to a non-conducting position (Yoder et al., 2018). A limitation of the available structures is the fact that the open structures of ASICs were not opened by acidic pH, but by gating-modifying toxins (Baconguis and Gouaux, 2012, Baconguis et al., 2014, Dawson et al., 2012). Since the opening was not induced by protonation, the conformation of other structural parts may differ from that of H^+^-opened ASICs, as discussed (Yoder and Gouaux, 2020).

### Structural interpretation of VCF with fluorophore-quencher pairing

VCF with fluorophore-quencher pairing indicates whether the residues of a given pair move towards, or away from each other during a defined transition. VCF does not indicate the distance change in absolute terms between the two partners, it does not indicate which of the two partners moves, and since fluorophores are large molecules, the actually detected distance change may occur at a position slightly besides the introduced Cys residue. A ΔF signal can also be caused by a reorientation of side chains. We have therefore applied a SMD-based strategy to evaluate the VCF-predicted distance changes, and to allow a structural interpretation. Below, only the VCF-predicted changes that were validated by the SMD simulations are considered.

### Conformational changes in the proximity of the β1-β2 linker during channel opening and desensitization

The VCF experiments, after validation by SMD simulations, indicate distance changes during activation between the β1-β2 linker and different structural elements of a neighboring subunit, as illustrated in Figure 8B: an approaching towards the lower end of the β9-α4 palm-thumb loop (’T289, β-turn) and the α5-β10 loop (’E359), and a distance increase to the close-by ‘E358, whose side chain moves away from Ser83. The VCF experiments highlight here fast distance changes between subunits, involving the thumb-palm loops (Figure 9A-B). The SMD analysis suggests that for the pairs of Ser83 with ‘Gln358 and ‘Glu359, mostly the thumb residues move, while for the Ser83-’Thr289 pair there is no evidence that one of the residues moves more than the other. These movements are followed by conformational changes predicted from the VCF kinetics to occur between opening and desensitization, and some correlated with desensitization (Figure 9C). Within the palm of each subunit, an approaching is observed between the β1-β2 linker and 1) Y417 of β12, and 2) Pro205 and Arg206 of the β5-β6 loop. This is accompanied by an approaching between the β1-β2 linker and ‘Asp357 as well as ‘Thr289, and a distance increase to ‘Gln358 and ‘L369 of a neighboring subunit. The generally slow conformational changes in the palm further support the tight link between the palm and desensitization, adding to the previously demonstrated importance of the β1-β2 linker in determining the desensitization kinetics (Coric et al., 2003), the established role of the β11-β12 linker with whom the β1-β2 linker tightly interacts (Springauf et al., 2011, Rook et al., 2020, Baconguis and Gouaux, 2012), as well as the importance of the lower palm β strands (Roy et al., 2013, Wu et al., 2019) in desensitization. The approaching of ‘D357 to β1-β2 linker residues may be related to slow conformational changes in the acidic pocket, which are not predicted from the structure comparison, for which there is however strong evidence from VCF experiments (Vullo et al., 2017).

### ASIC pH sensing and signaling towards the gate

Mutations of titratable residues in the palm, wrist and acidic pocket affect the pH dependence of ASICs, and although crystal structures predict an important role of the acidic pocket in ASIC activation (Jasti et al., 2007), it was shown that pH sensing in the acidic pocket is not required for the generation of transient H^+^-induced ASIC currents (Vullo et al., 2017). In the lower palm, putative pH-sensing residues were identified in several β strands (Liechti et al., 2010, Krauson et al., 2013). Mutation of the wrist residue His73 to Ala shifted the pH dependence of activation to more acidic values, and simultaneous mutation of His72 and His73 to Asn suppressed pH-induced currents completely (Paukert et al., 2008).

We show here that activation-related conformational changes occur in the finger and in the acidic pocket. Such conformational changes can likely be transmitted down to the pore from the finger and acidic pocket via the thumb and the thumb-palm loops to the TM1 involving the interactions between aromatic residues of the TM1 and the β-turn, and/or via the central scaffold and subunit interactions to the palm and from there to the transmembrane domains (Figure 9A). Protonation events in the palm and wrist would affect the transmission of conformational changes initiated farther up, or may drive in part the large conformational changes of these domains, which would - due to their proximity - also change the conformation of the transmembrane domains or affect the position of the β-turns. Mutation of the interacting residues between the β-turn and the TM1 disrupts ASIC1a currents (Li et al., 2009), leading to a lowering of the cell surface expression, and rendering the cell surface-resident channels non-functional (Jing et al., 2011). Normal mode analysis had suggested a high correlation between movements of the β-turn and the upper TM1 (Yang et al., 2009), highlighting together with the experimental data a possible importance of the β-turn for ASIC activation.

In conclusion, we show here that upon extracellular acidification, fast conformational changes occur simultaneously in different ASIC domains, consistent with the existence of multiple protonation sites. Analysis of the kinetics and the direction of conformational changes in the palm and palm-thumb loops highlights rapid events in the palm-thumb loops, which may therefore constitute a preferred communication pathway between protonation sites and the channel gate in the context of activation.

## Materials and Methods

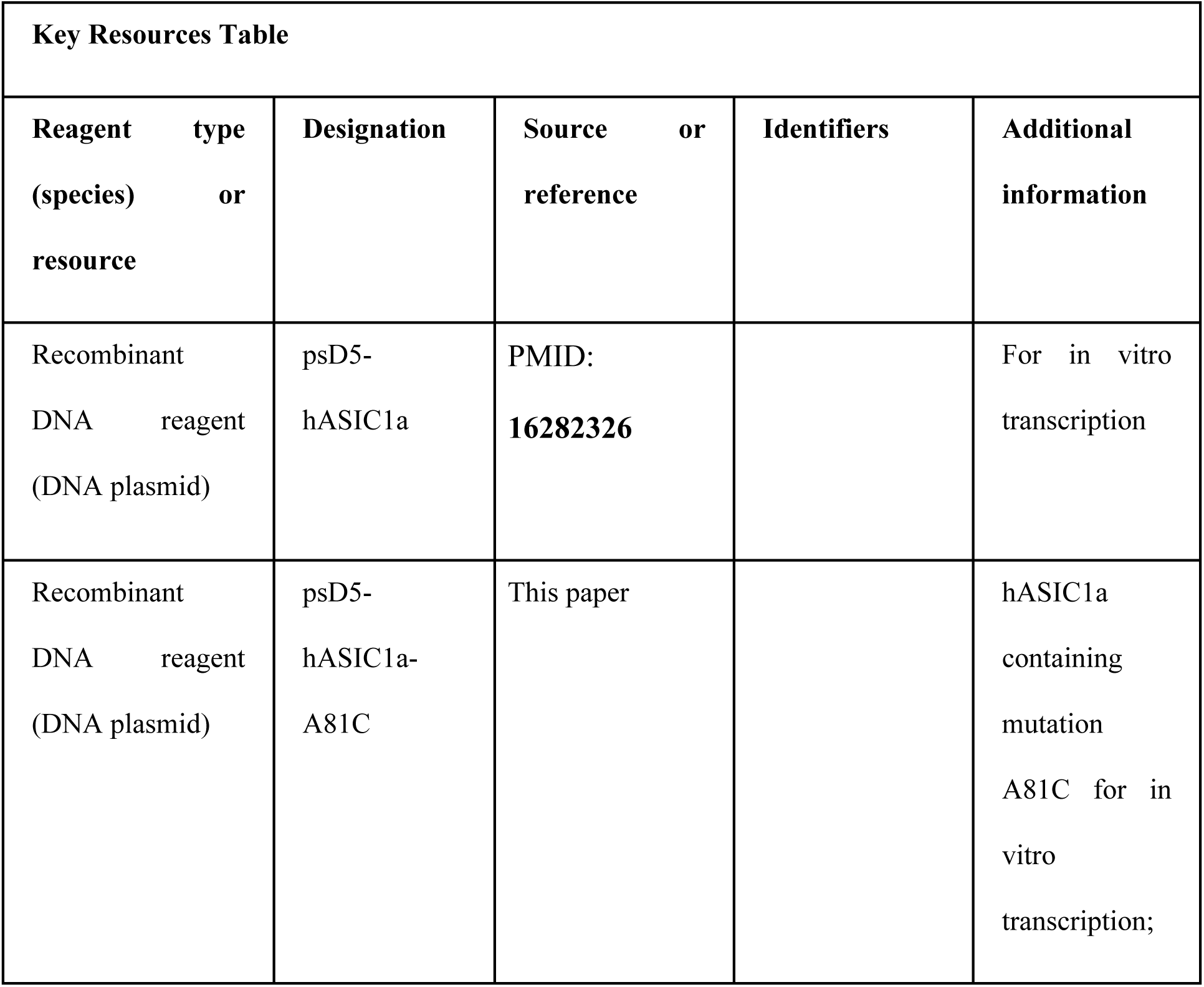
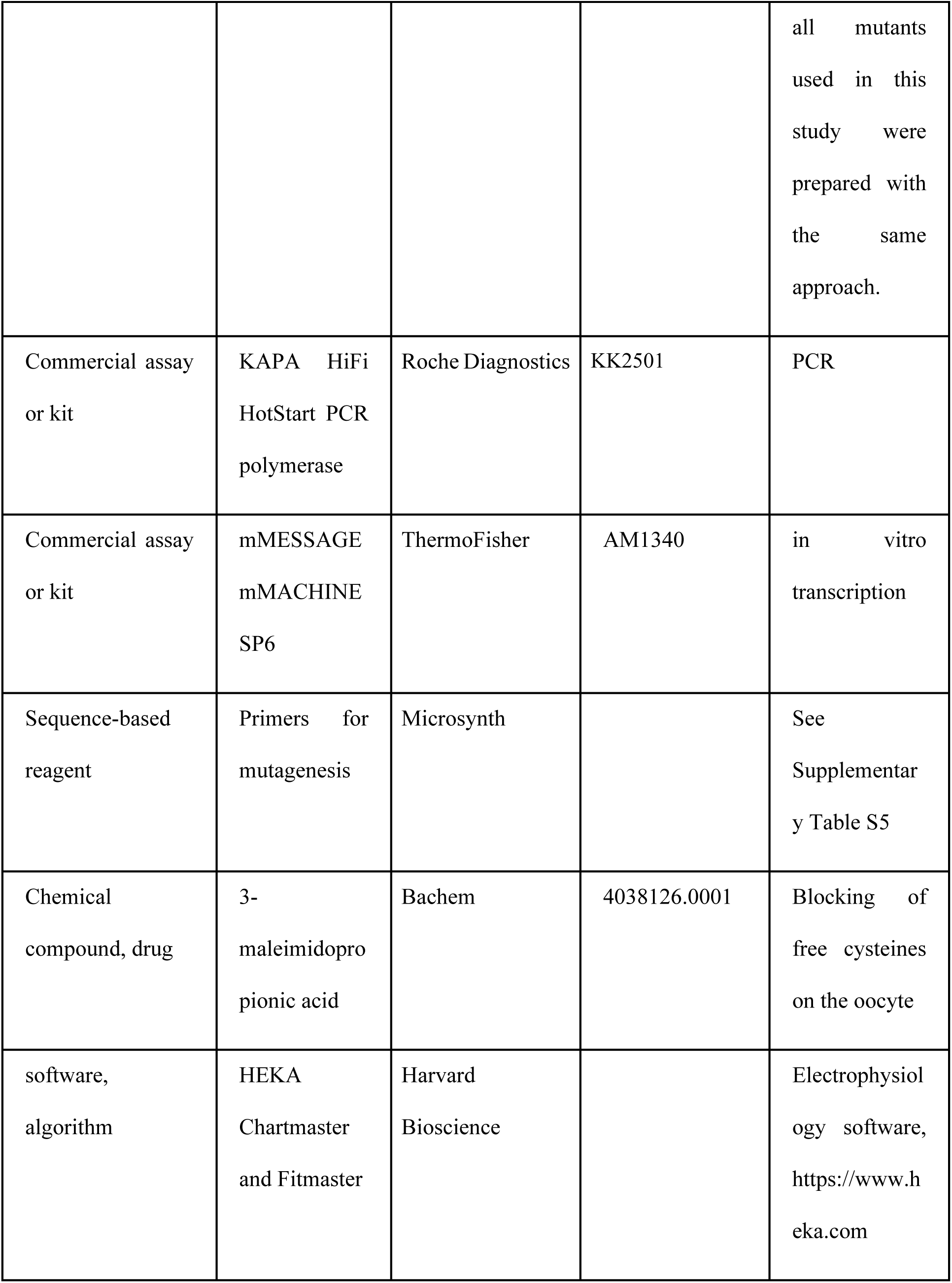
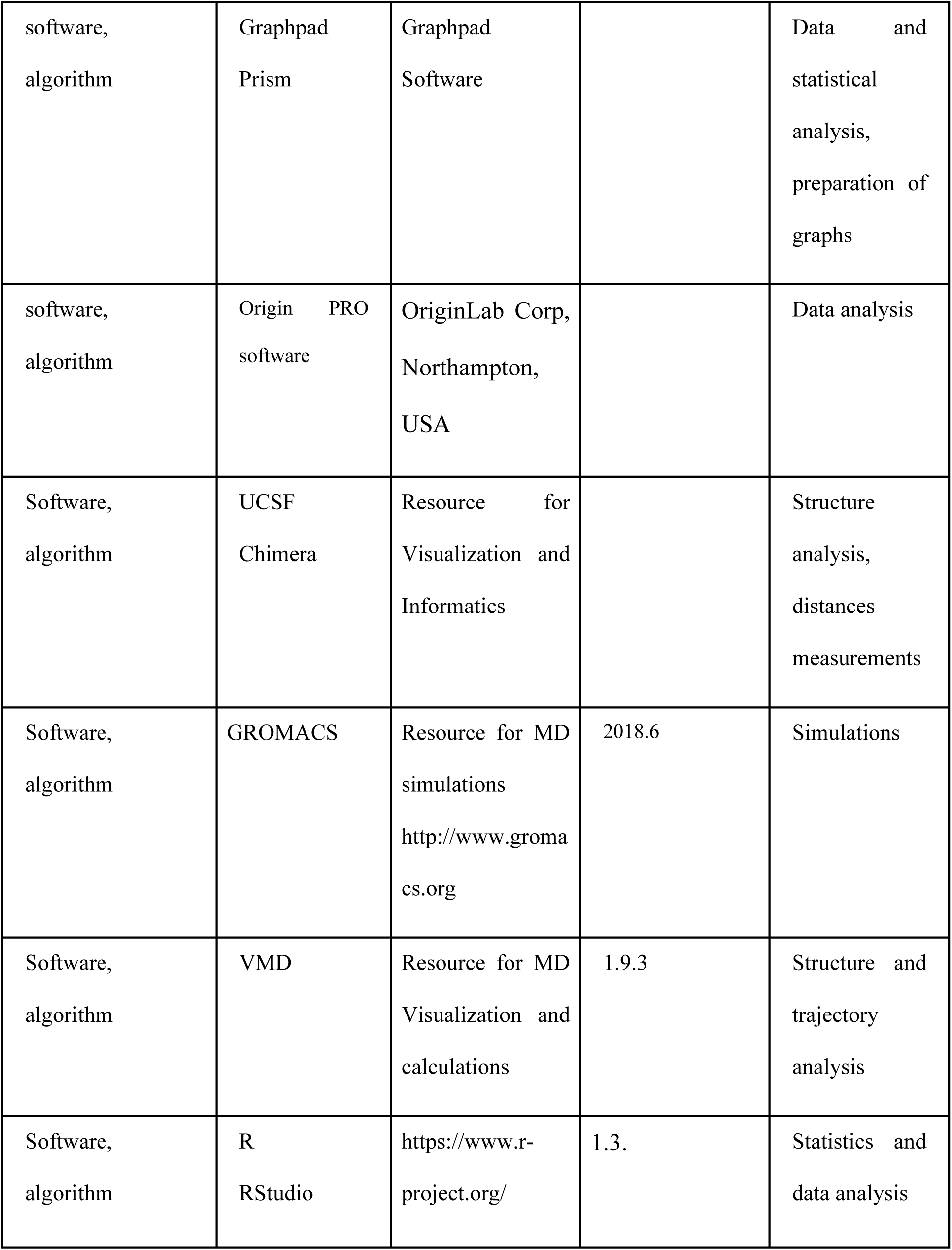
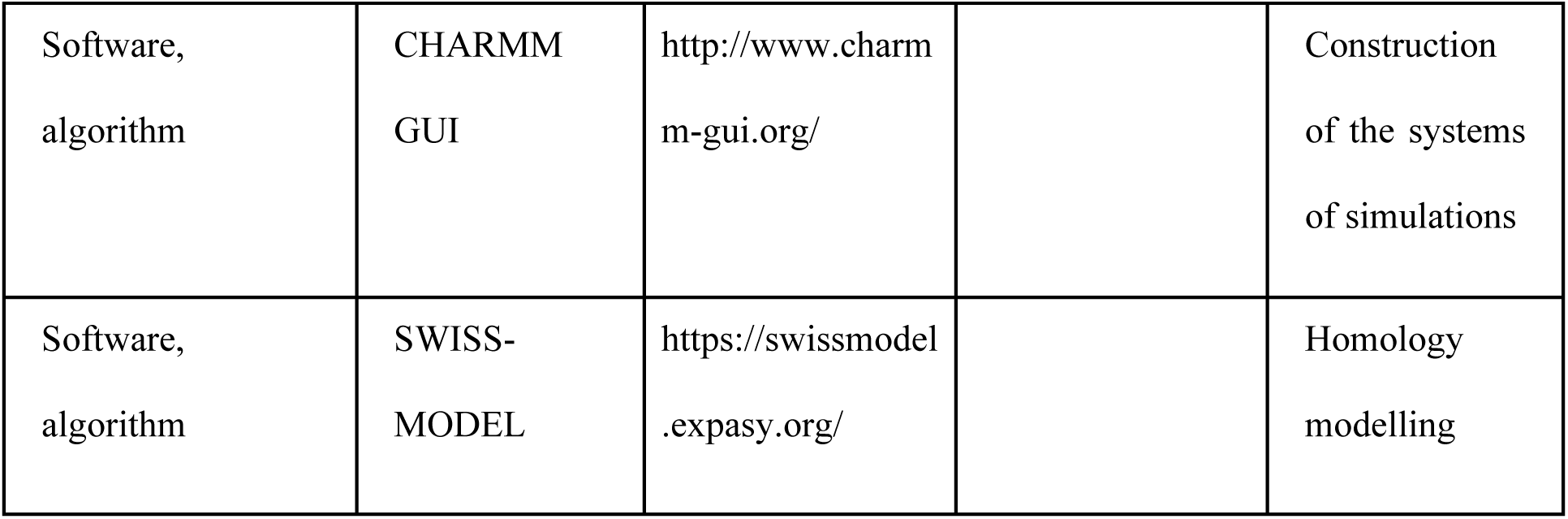

| Key Resources Table |  |  |  |  |
| --- | --- | --- | --- | --- |
| Reagent type<br>(species) or<br>resource | Designation | Source or<br>reference | Identifiers | Additional<br>information |
| Recombinant<br>DNA reagent<br>(DNA plasmid) | psD5-hASIC1a | PMID:<br>**16282326** |  | For in vitro<br>transcription |
| Recombinant<br>DNA reagent<br>(DNA plasmid) | psD5-hASIC1a-A81C | This paper |  | hASIC1a<br>containing<br>mutation<br>A81C for in<br>vitro<br>transcription; |
|  |  |  |  | all mutants used in this study were prepared with the same approach. |
| Commercial assay or kit | KAPA HiFi HotStart PCR polymerase | Roche Diagnostics | KK2501 | PCR |
| Commercial assay or kit | mMESSAGE mMACHINE SP6 | ThermoFisher | AM1340 | in vitro transcription |
| Sequence-based reagent | Primers for mutagenesis | Microsynth |  | See Supplementar y Table S5 |
| Chemical compound, drug | 3-maleimidopropionic acid | Bachem | 4038126.0001 | Blocking of free cysteines on the oocyte |
| software, algorithm | HEKA Chartmaster and Fitmaster | Harvard Bioscience |  | Electrophysiology software, https://www.heka.com |
| software,<br>algorithm | Graphpad<br>Prism | Graphpad<br>Software |  | Data and<br>statistical<br>analysis,<br>preparation of<br>graphs |
| software,<br>algorithm | Origin PRO<br>software | OriginLab Corp,<br>Northampton,<br>USA |  | Data analysis |
| Software,<br>algorithm | UCSF<br>Chimera | Resource for<br>Visualization and<br>Informatics |  | Structure<br>analysis,<br>distances<br>measurements |
| Software,<br>algorithm | GROMACS | Resource for MD<br>simulations<br>http://www.gromacs.org | 2018.6 | Simulations |
| Software,<br>algorithm | VMD | Resource for MD<br>Visualization and<br>calculations | 1.9.3 | Structure and<br>trajectory<br>analysis |
| Software,<br>algorithm | R<br>RStudio | https://www.r-project.org/ | 1.3. | Statistics and<br>data analysis |
| Software,<br>algorithm | CHARMM<br>GUI | http://www.charmm-gui.org/ |  | Construction<br>of the systems<br>of simulations |
| Software,<br>algorithm | SWISS-MODEL | https://swissmodel.expasy.org/ |  | Homology<br>modelling |

### Molecular biology

The human ASIC1a cDNA construct (Garcia-Anoveros et al., 1997) was cloned into a pSP65 vector containing 5’- and 3’-untranslated sequences for expression in *Xenopous* oocytes. As reported recently, this construct contains the mutation G212D (Vaithia et al., 2019). Point mutations were introduced by site-directed mutagenesis using KAPA HiFi HotStart PCR polymerase (Roche Diagnostics, Rotkreuz, Switzerland). All mutations were verified by sequencing (Synergen Biotech). Complementary RNAs were synthetized *in vitro* using the mMESSAGE mMACHINE SP6 kit (Thermofisher).

### Oocyte expression

Surgical removal of oocytes was carried out as described previously (Liechti et al., 2010). Healthy oocytes of stages V and VI were collected from adult female *Xenopous laevis* in accordance with the Swiss federal law on animal welfare and approved by the committee on animal experimentation of the Canton de Vaud. Oocytes were injected with 1-50 ng of cRNA encoding hASIC1a WT and mutants. Oocytes used for VCF experiments were incubated after cRNA injection for 1h in Modified Barth’s Solution (MBS) containing 10 mM 3-maleimidopropionic acid (Bachem) to modify free cysteine residues of proteins natively expressed on the cell membrane, and then maintained at 19°C in MBS, composed of (mM): 85 NaCl, 1 KCl, 2.4 NaHCO_3_, 0.33 Ca(NO_3_)_2_, 0.82 MgSO_4_, 0.41 CaCl_2_, 10 HEPES and 4.08 NaOH.

### Electrophysiology

Two-electrode voltage clamp (TEVC) and voltage-clamp fluorometry (VCF) experiments were conducted at room temperature (20-25°C) 1-2 days after cRNA injection. All oocytes used for TEVC and VCF experiments had been previously labelled in the dark with 5 μM CF488 maleimide (Biotium) or AlexaFluor488 maleimide (Invitrogen) at room temperature for 15 min. Oocytes were placed in a RC-26Z recording chamber (Warner Instruments) and impaled with glass electrodes filled with 1M KCl, and continuously perfused by gravity at a rate of 5-15mL/min. A specially designed chamber was used to measure the kinetics of the fluorescence changes (ΔF) and current from about the same oocytes surface, as described previously (Vullo et al., 2017). In this chamber, the solution flows under the oocyte (Figure 1C). Macroscopic currents were measured at a holding potential of −40 mV with a TEV-200A amplifier (Dagan Corporation). Data were recorded with Chartmaster software (HEKA Electronics) at a sampling rate of 1 ms. Standard recording solutions contained (mM): 110 NaCl, 2 CaCl_2_, and 10 HEPES for pH ≥ 6.8. For solutions with a pH < 6.8, HEPES was replaced by 10 mM MES. The pH was adjusted using NaOH and HCl. For the current and ΔF measurements, channels were stimulated once every minute for 10s (20s for pH dependence measurements of ΔF).

The speed of solution change in the area used for the measurement of the fluorescence and the current signal of the chamber used for the kinetic measurements was determined by the same approach as described previously for a different chamber (Bonifacio et al., 2014). The constitutively active, Na^+^-selective ENaC was expressed in oocytes, and oocytes were labeled before the experiment with the membrane-impermeable, pH-sensitive fluorophore 5(6) FAM SE [5-(and-6)-Carboxyfluorescein, succinimidyl ester] mixed isomers (Biotium, Chemie Brunschwig, Basel, Switzerland). In the experiments, the solution was changed from one containing K^+^ as cation, at pH7.4 to one in which K^+^ was replaced by Na^+^, and whose pH was 6.0. This allowed the simultaneous determination of the solution change by fluorescence and by current measurement without the involvement of a channel activation, since ENaC is constitutively active, but not permeable to K^+^. The rise time (RT) of the solution change was 355 ± 37 ms (current) and 296 ± 34 ms (ΔF), and the ratio RTΔF/RTI was 0.85 ± 0.07 (n=8). These kinetics are slower than the current and ΔF measured for ASIC activation by pH in many experiments. It is known from experiments with ultrarapid perfusion systems and excised patches that the intrinsic kinetics of ASIC activation are of the order of ∼10 ms (Bassler et al., 2001, Sutherland et al., 2001, Alijevic et al., 2020, Vaithia et al., 2019). Therefore, the ASIC activation kinetics are in most studies limited by the speed of the perfusion change. Our control experiments indicate that in experiments with ASICs, the current and ΔF maximum was reached before the solution change was complete, indicating that the peak was reached at a lower H^+^ concentration than the one of the acidic solutions. Therefore, the actual acidic pH of the kinetic measurements is more alkaline than the pH of the acidic solution. Since in our experiments, relative differences between current and ΔF kinetics, measured simultaneously, are analyzed, the conclusions about the speed of the ΔF relative to the current signal, and with this the attribution to a given functional transition, is valid. It needs however to be taken into account that the absolute speed of these kinetics is limited by the kinetics of the solution change, and that the actual pH of the measurement of the kinetics is slightly more alkaline than that of the test solution.

### Fluorescence measurements

The VCF setup was equipped with an Intensilight mercury lamp (C-HGFI; Nikon) and a 40x Nikon oil-immersion objective to detect the fluorescence signal emitted by fluorophore-labelled oocytes. The optical signal was then converted into current units by a photodiode (S1336-18BQ; Hamamatsu Photonics) coupled to the head stage of an amplifier (List-EPC-7; HEKA). A low-pass eight-pole Bessel filter was used to amplify and filter the signal at 50 Hz. Changes in fluorescence intensity (ΔF) were normalized to the total fluorescence signal (F). Specificity of the fluorescence signals was assessed by exposing the oocytes to a slightly acidic pH (pH6.7) for 50s, which puts the channels in the desensitized state, before they were stimulated with pH6. This protocol did not generate ionic current, because the channels were desensitized before the acidification to pH6. If a Cys mutant showed a substantial fluorescence signal after application of this protocol, the signal was considered as potentially non-specific. It can however not be excluded that such a signal may be due to a transition between different sub-states, as for example two desensitized states. As a measure of the specificity of the signal (or a component of the signal), the ratio of the ΔF/F induced by acidification from the desensitized relative to the ΔF/F induced by acidification from the closed state was calculated (Table S3). The lower the ratio, the higher is the confidence that the fluorescence signal is specific.

### Kinetic model, data analysis and statistics

To predict the time dependence of the ASIC1a probability of being in a given functional state, ASIC1a WT was modeled according to a previously published kinetic model that is based on the Hodgkin-Huxley formalism, containing an activation and a sensitization gate, and containing 4 functional states, closed, open, closed-desensitized and open-desensitized (Alijevic et al., 2020). Only the open state conducts current. The same model was also used as a basis for the simulation of ΔF traces (see below).

Experimental data were analyzed with the software Fitmaster (HEKA Electronics) and with Origin PRO software (OriginLab Corp, Northampton, USA). pH-response curves were fit to the Hill function: [(I = I_max_/(1+ (10^-pH^ /10^-pH^)^nH^], where I_max_ is the maximal current, pH_50_ is the pH that induces 50% of the maximal current amplitude, and nH is the Hill coefficient. Steady-state desensitization (SSD) curves were fitted with an analogous equation.

The results are presented as mean ± SEM. They represent the mean of n independent experiments on different oocytes. Statistical analysis was done with t-test where two conditions were compared, or with One-way ANOVA followed by Dunnett’s or Tukey multiple comparisons test for comparison of >2 conditions for normally distributed data, and Kruskal-Wallis and Dunn’s test for non-normally distributed data (Graphpad Prism 8). ΔF and current kinetics were considered as correlated for a given mutant and pH condition if the steepness of the linear regression of a plot of the rise time (as time to pass from 10 to 90% of the full amplitude) of ΔF as a function of the rise time of the current kinetics was between 0.75 and 1.33. Structural images were generated with the UCSF Chimera software (Pettersen et al., 2004).

### Molecular Dynamics simulations

Homology models of human ASIC1a were constructed from chicken ASIC1a which shares 90% identity with its human homolog, using structures representing the closed (PDB code 5WKU), open (4NTW) and desensitized (4NYK) structures (Gonzales et al., 2009, Baconguis et al., 2014, Yoder and Gouaux, 2018, Yoder et al., 2018) with SWISS-MODEL (Biasini et al., 2014). The molecular systems were constructed as done previously (Bignucolo et al., 2020). They contained ∼250 or ∼220 1-palmitoyl-2-oleoyl-sn-glycero-3-phosphocholine (POPC) molecules in each leaflet for the closed or open state, respectively, and the total number of atoms was 281’000 and 244’000. The GROMACS package, version 2018.6, was used to conduct simulations with the CHARMM force-field (MacKerell et al., 1998), versions v27 for proteins (Mackerell et al., 2004) and v36 for lipids (Klauda et al., 2010). The systems were equilibrated following the CHARMM-GUI protocol (Jo et al., 2007).

The parameters for the steered molecular dynamics (SMD) were set as done previously (Bignucolo and Berneche, 2020), with the difference that here the constant force was replaced by a harmonic potential without initial velocity and with an initial force of 30 kJ/mol/Å. For each pair of residues, a target distance was defined as a function of the VCF-predicted distance change. The target distance was calculated by adding 5Å to or subtracting 5Å, respectively, from the distance measured between the two atoms before the transition, depending on whether VCF predicted an increase or a decrease of the distance. For example, the distance between the side chains of residues 81 and 417 is ∼ 8.6 Å in the open state homology model after equilibration. The corresponding VCF signal was negative, indicating an approaching between the fluorophore at position 81 and Tyr417. Thus, the target distance between these two residues was set to 8.6 – 5 = 3.6Å. The kinetics of the VCF signal indicated that it occurs during the open-desensitized transition. The initial and target distances are reported in the table S4. For simulations starting from the closed state, the forces were exerted on the center of mass of the most distal heavy atoms of the side chain (Ser83 and Thr289: OG; Leu207 and Leu369: CD1, CD2; Gln358: OE1, NE2; Glu359: OE1, OE2) whereas for simulations starting from the open state, where a total of 42 restraints were applied, the forces were exerted on the Cβ atoms. Three independent simulations of 140 and 170 ns duration were conducted for the closed-open and open-desensitized transitions, respectively, so that for each transition nine different residue pairs could be investigated. The main purpose of these simulations was to rank the movements deduced from the VCF experiment in terms of structural plausibility and to interpret them in structural terms. It was assumed that a VCF-deduced movement that corresponds to a native conformational change, or at least that does not require excessive conditions, e.g. an excessive temperature, would be approached easily and would require a rather weak pull force. Since a harmonic potential implies that the intensity of the force and the deviation from the target value are correlated, a rapid approach of the target distance together with a decrease of the exerted force indicates a low energy barrier. For the analysis, the period after the first 100 ns of simulations was used. The simulations for the closed-open transition included constraints from 5 residue pairs, while the simulations for the open-desensitized transition included constraints from 15 residue pairs (Tables 2 and S4). Whereas the achievable distances are largely defined by geometrical considerations, forces exerted on neighbouring residues can add to each other (see disposition of the arrows in Figure 7-figure supplement 1). In order to compensate qualitatively to this tendency, the force threshold was reduced in the open-desensitized transition analysis, because it harbours much more distance constraints than the closed-open transition analysis. The distance requirements were considered validated by the SMD if the deviation from the target distance was lower than 2.2 Å and the applied force, in absolute value, was lower than 6 kJ/mol/Å for the closed-open transition. The threshold for the applied force was set to. 5 kJ/mol/Å for the open-desensitized transition. Table 2 reports the calculated deviations and applied forces, as well as the validation output for both investigated transitions. Data were analysed using R (https://www.R-project.org/), python, VMD (Humphrey et al., 1996) (https://www.ks.uiuc.edu/) and with tool common language (tcl) in-house scripts. The Tukey Honest Significant Differences test was performed in case of ANOVA with multiple factor levels (Yandell, 1997). The timeline plugin of VMD was used to extract the displacement values.

### Simulation of ΔF signals from a kinetic ASIC1a model

As one possible interpretation of the ΔF signals it was assumed that a certain degree of fluorescence was associated with each functional state (Figure 1F-G). To this end, the kinetic model describing the function of ASIC1a WT, described in the *Methods* (Alijevic et al., 2020), was used, and scaling factors between −1 and +1 relating the fluorescence to each of the functional states (C, closed; O, open; CD, closed-desensitized; OD, open-desensitized) were chosen to reproduce best the measured ΔF traces. Specifically, ΔF was modeled as

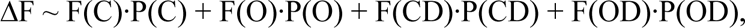

where the symbol “∼” denotes proportionality, P(C), P(O), P(CD) and P(OD) are the probabilities of the channel to be in the corresponding states (which evolve with time but always sum up to 1) and F(C), F(O), F(CD) and F(OD) are the corresponding constant scaling factors. The traces in Figure 1G show modeled ΔF traces for the case that ΔF was associated to one single functional state, or associated to either the activation or sensitization gate. The model assumes that fluorescence can be positively or negatively associated with any of the four states. We aimed to use the simplest model (least number of non-zero factors) that described the measured ΔF sufficiently well. For ease of interpretation, we included an F(C) different from 0 only if the trace could not be described with attributing values to F(C), F(O) and/or F(CD) alone. For the large majority of ΔF traces it was possible to find combinations of scaling factors that reproduced the experimental trace reasonably well (see for example Figure 1D). The values of F(C), F(O), F(CD) and F(OD) describing each model are noted in blue below the simulated traces, and are presented in Supplementary Table S2. These factors were estimated empirically as follows. For some recurring, relatively simple patterns of the ΔF signal, defined rules were used to derive the factors. Simple transient signals containing in addition a sustained component (e.g. E63C) were characterized by a F(O) and F(OD). F(O) was set equal to −1 or +1 depending on the polarity of the ΔF signal, and F(OD) was chosen to reproduce the quantitatively determined ΔFsust/ΔFpeak ratio. Many of the measured ΔF signals were sustained, without transient component (e.g. K105C). Simulations with the kinetic model showed that the kinetics of the ΔF onset depend almost exclusively on F(O) and F(OD). The absolute value of F(OD), (abs(F(OD))) needs to be > abs(F(O)) to ensure that there is no transient component. F(OD) was set to −1 or 1 in these cases, and F(O) was based on experimentally determined kinetics at pH6.0 of the RT of the fluorescence onset (RTF) relative to the current rise time (current activation, RTAI) and the current decay time (current desensitization, RTDI) as (RTF-RTAI)/(RTDI-RTAI). This ratio was also determined with the kinetic model for a number of values of F(O), and a function relating F(O) to the (RTF-RTAI)/(RTDI-RTAI) ratio was derived from the model, and used to calculate F(O) for each mutant showing a simple sustained signal (Supplementary Table S2, Model parameters).

For acidification from pH7.4 to pH6.0, the P(CD) is ∼0 during the acidification and increases immediately upon returning to pH7.4, and decreases then slowly. This slow decrease is related to the recovery from desensitization. A slow decay of the ΔF signal upon returning to pH7.4 was observed in many mutants, as e.g. S83C (Figure 4B). In some cases, only a part of the decay is slow (I428C, Figure 2B). If abs(F(CD)) > abs(F(OD)), a partially sustained ΔF signal increases transiently upon returning to pH7.4 (e.g. Y71C, Figure 2B). F(O) and F(OD) do not influence the “off” kinetics of the ΔF signals. To obtain values of F(CD) for each mutant, the experimental “off” kinetics of the ΔF signal of each mutant were measured for the pH6.0-pH7.4 transition (RTFoff, Table S1). These kinetics were also determined for a model with F(OD)=1 and different values of F(CD), and the relationship in the model between F(CD) and RTFoff was determined. If abs(F(OD)) < 1, the F(CD)/F(OD) ratio determined the “off” kinetics (i.e. the RTFoff in the model F(OD)=1, F(CD)=1 was equal to that of F(OD)=0.5, F(CD)=0.5). The relationship between F(CD) and RTFoff determined in the model was then used to calculate F(CD) from the experimental RTFoff for each mutant (Supplementary Table S3, Model parameters).

## Acknowledgements

The authors thank Ivan Gautschi for expert experimental work, and Anand Vaithia and Zhong Peng for their comments on a previous version of the manuscript. This work was supported by the Swiss National Science Foundation grant 31003A_172968 to SK. This work was supported by grants from the Swiss National Supercomputing Centre (CSCS) under project s1037.

---

**Figure 1-figure supplement 1.**
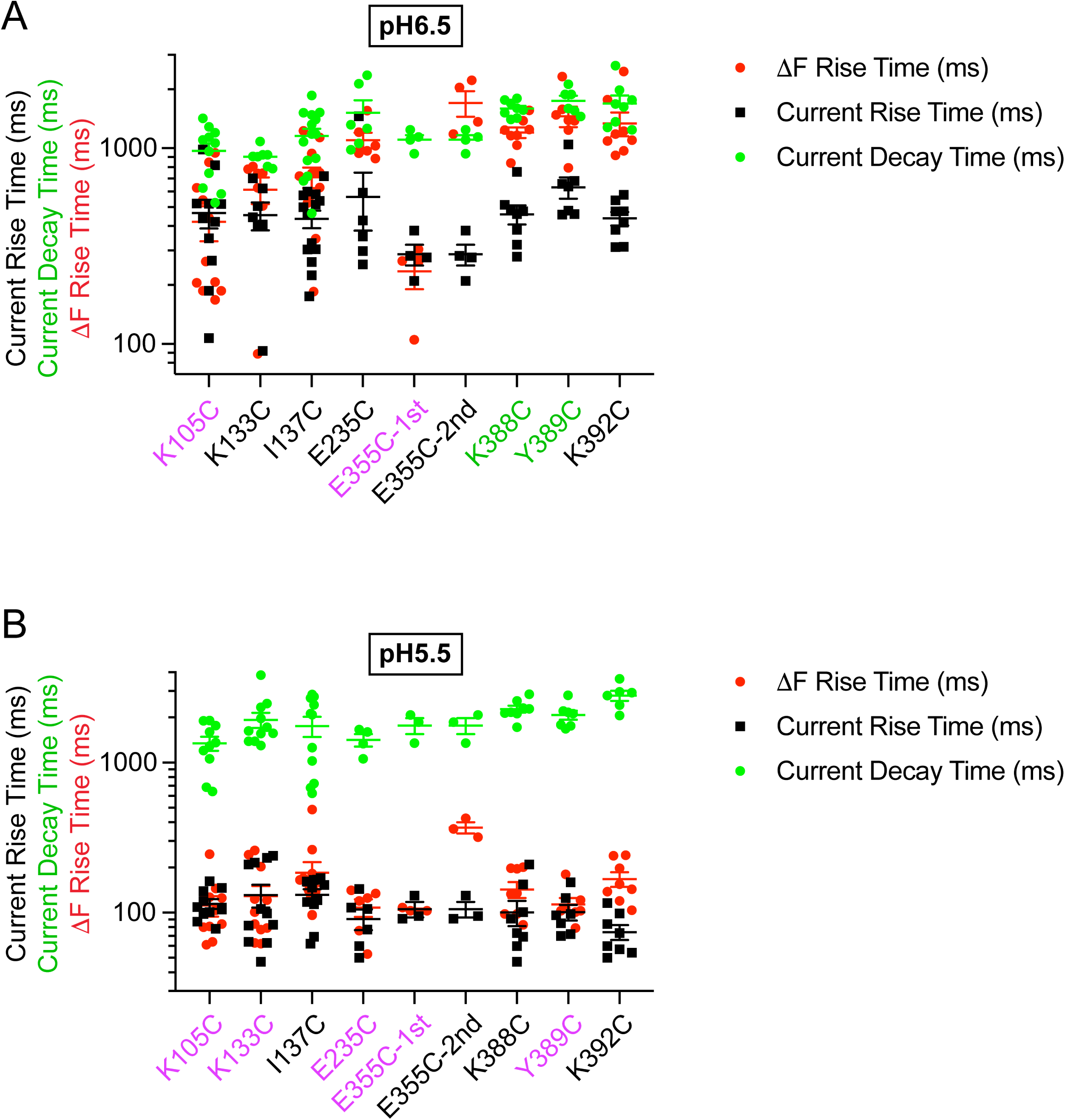
Kinetics of mutants with rapid ΔF located outside the wrist at pH6.5 and 5.5. Current RT and decay time and ΔF RT obtained at pH6.5 (**A**) or 5.5 (**B**), as indicated, n=3-14. The color of the labels of the mutants indicates that the ΔF onset kinetics are correlated with current appearance (purple) or with current desensitization (green; *Materials and Methods*).

**Figure 2-figure supplement 1.**
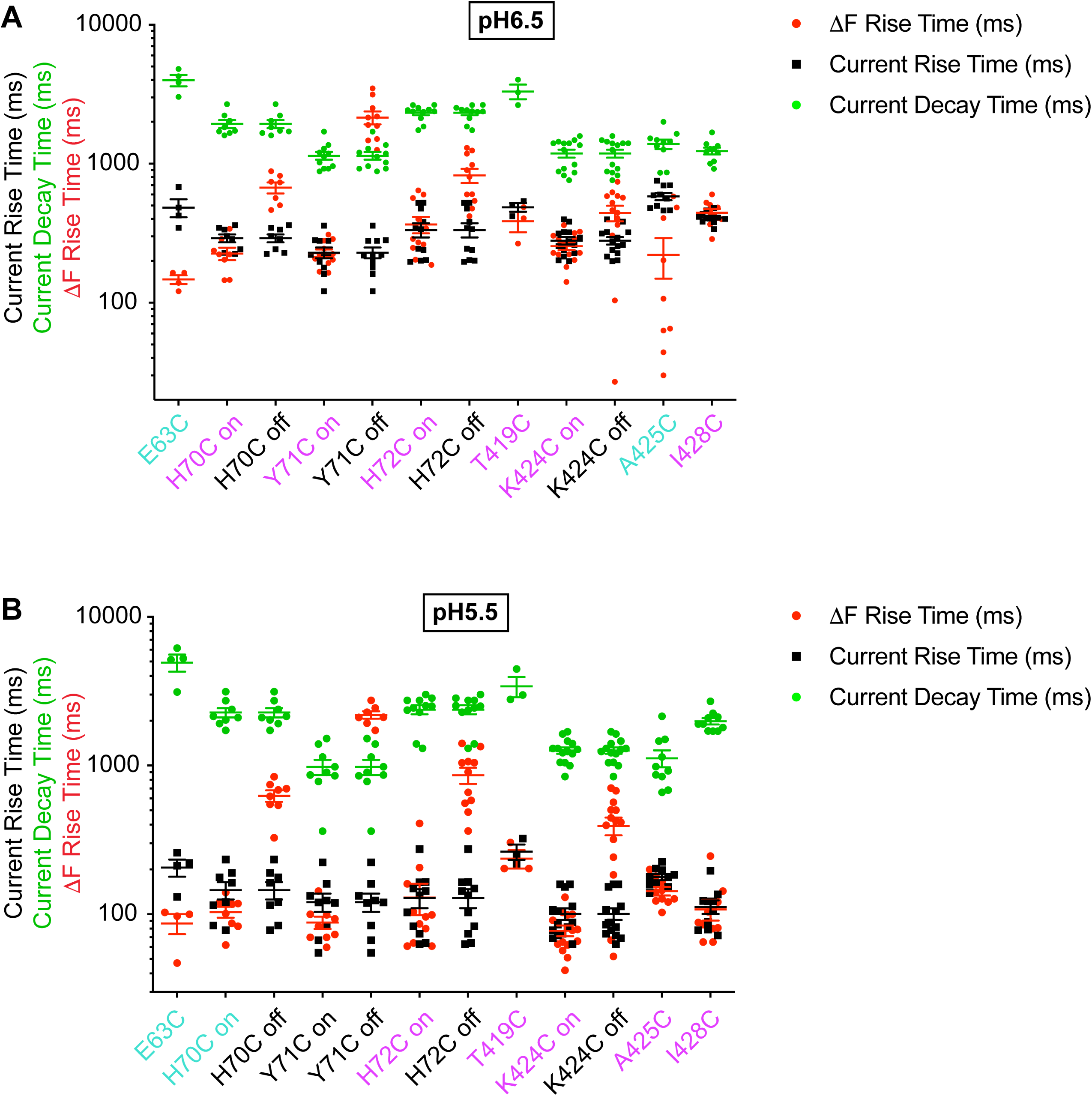
Rapid ΔF kinetics of wrist mutants at pH5.5 and 6.5. Current RT and decay time and ΔF RT obtained at pH6.5 (**A**) or 5.5 (**B**), as indicated, n=3-14. The color of the labels of the mutants indicates that the ΔF onset kinetics are correlated with current appearance (purple) or are faster than current appearance (cyan; *Materials and Methods*).

**Figure 2-figure supplement 2.**
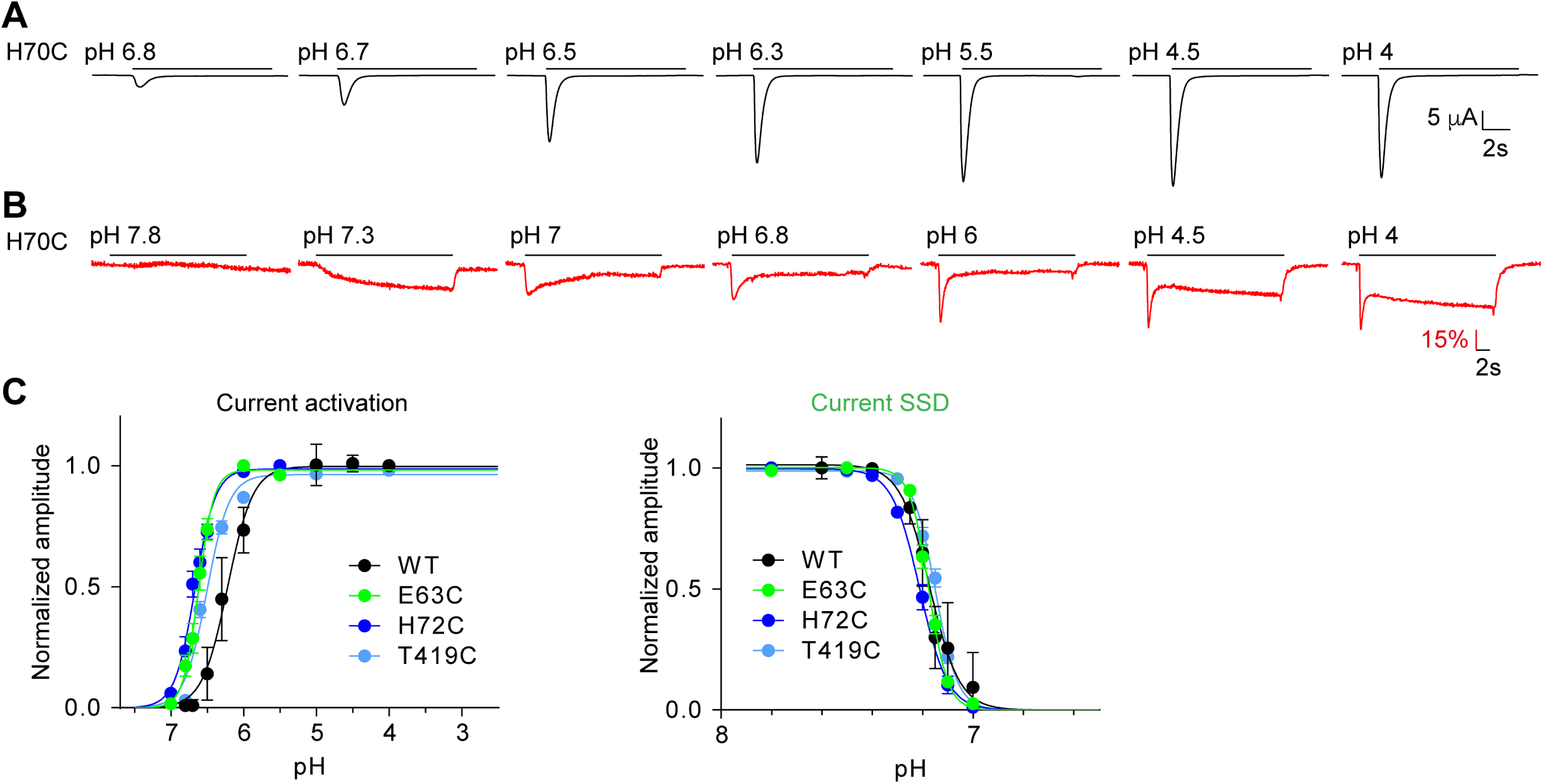
pH dependence of mutants involved in rapid conformational changes. **(A)** and **(B)** Representative traces of experiments measuring the pH dependence of the current and of ΔF, respectively of the mutant H70C. (**C**) Activation curves for the current and steady-state desensitization (SSD) curve of the indicated mutants. The pH protocols were as illustrated in (**A**) and (**B**) and described in *Materials and Methods*. For the analysis of the SSD pH dependence, the protocol was as follows: After exposure during 50s to the conditioning pH, the channels were exposed during 10s to pH4.5. This protocol was repeated at different conditioning pH values, and the normalized current is plotted as a function of the conditioning pH.

**Figure 3-figure supplement 1.**
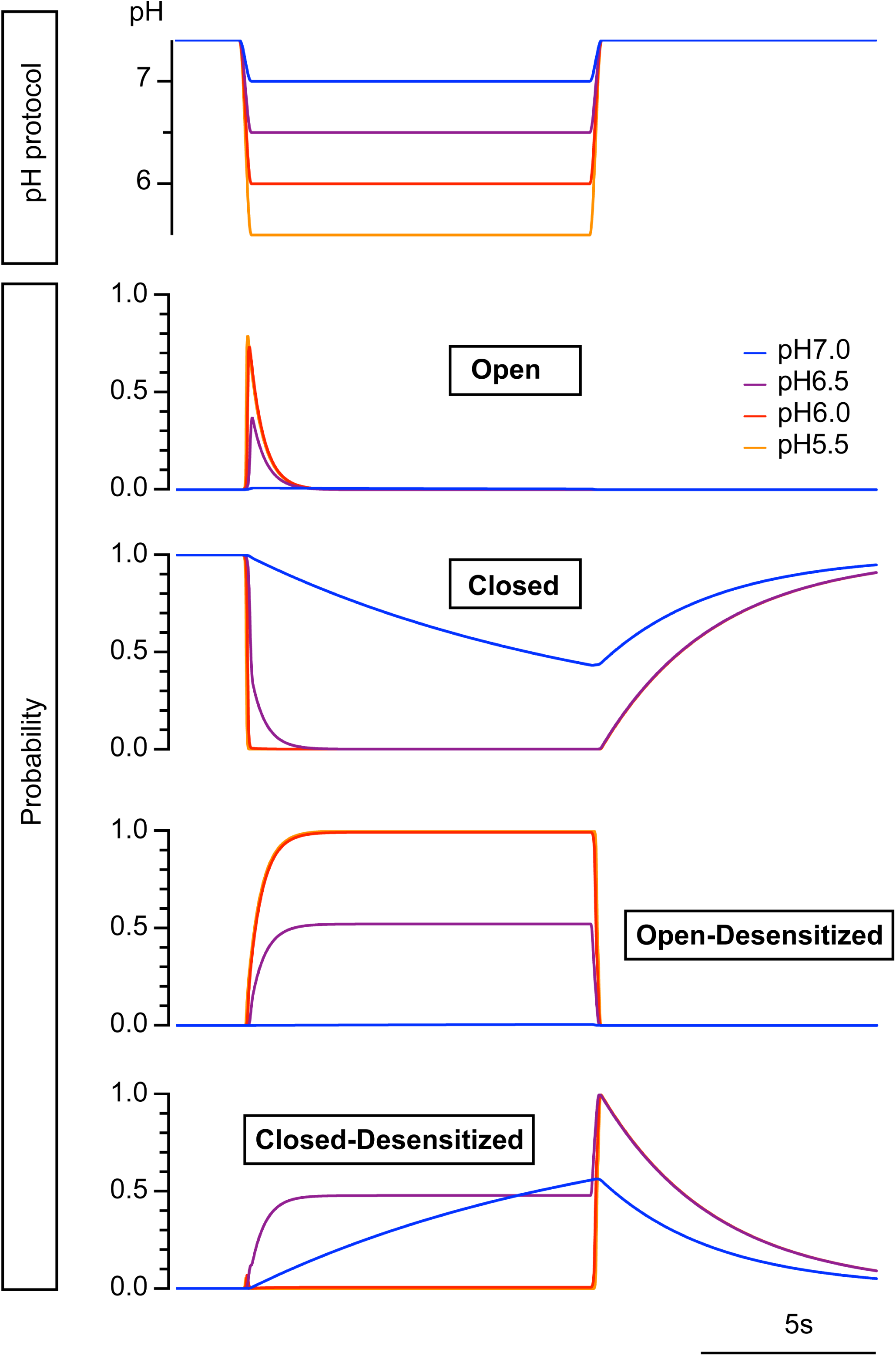
Modeled state probability at different pH conditions. Time course of probability of the four model states for a pH change from 7.4 to 7.0, 6.5, 6.0 and 5.5. The conditioning pH7.4 was changed to the acidic pH at 2s for a duration of 10s. The fall time of the pH change was set to 200 ms.

**Figure 4-figure supplement 1.**
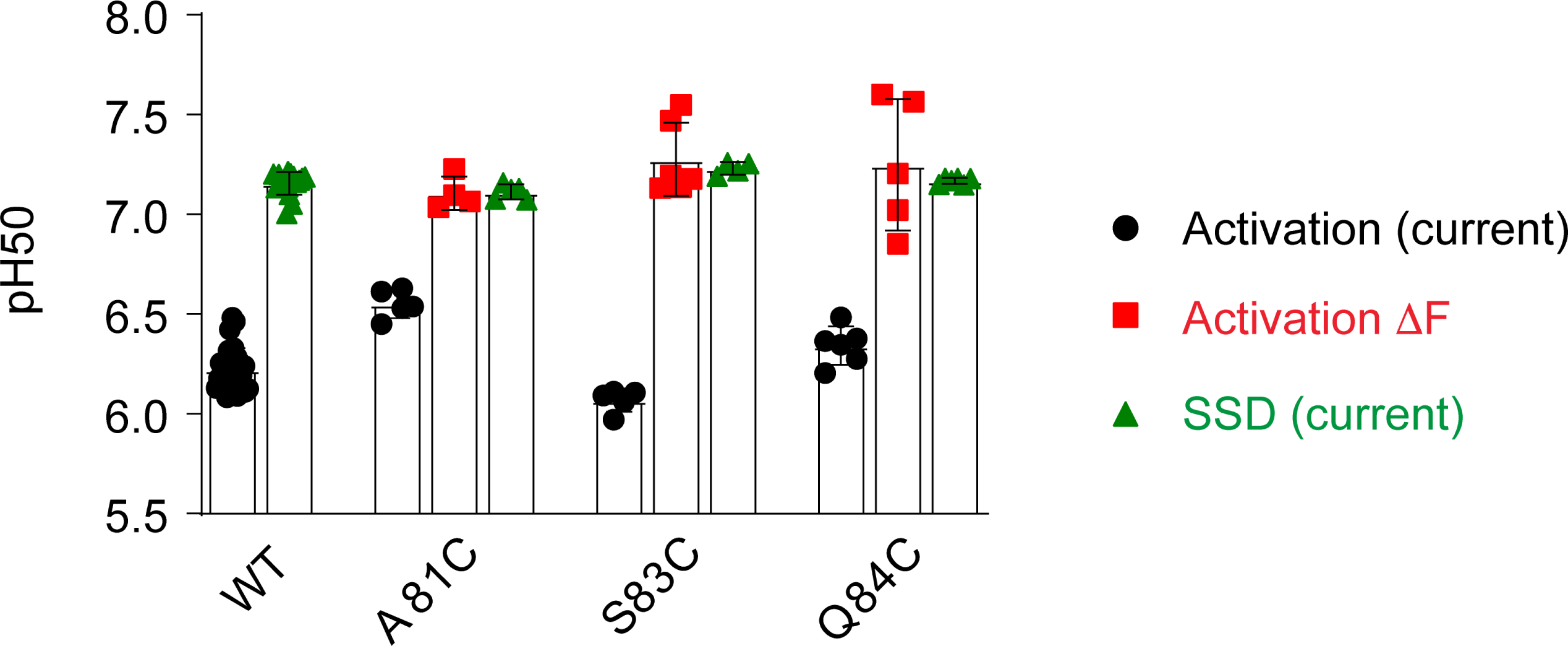
pH dependence of palm mutants. pH_50_ values of current activation and SSD, and of ΔF are indicated.

**Figure 5-figure supplement 1.**
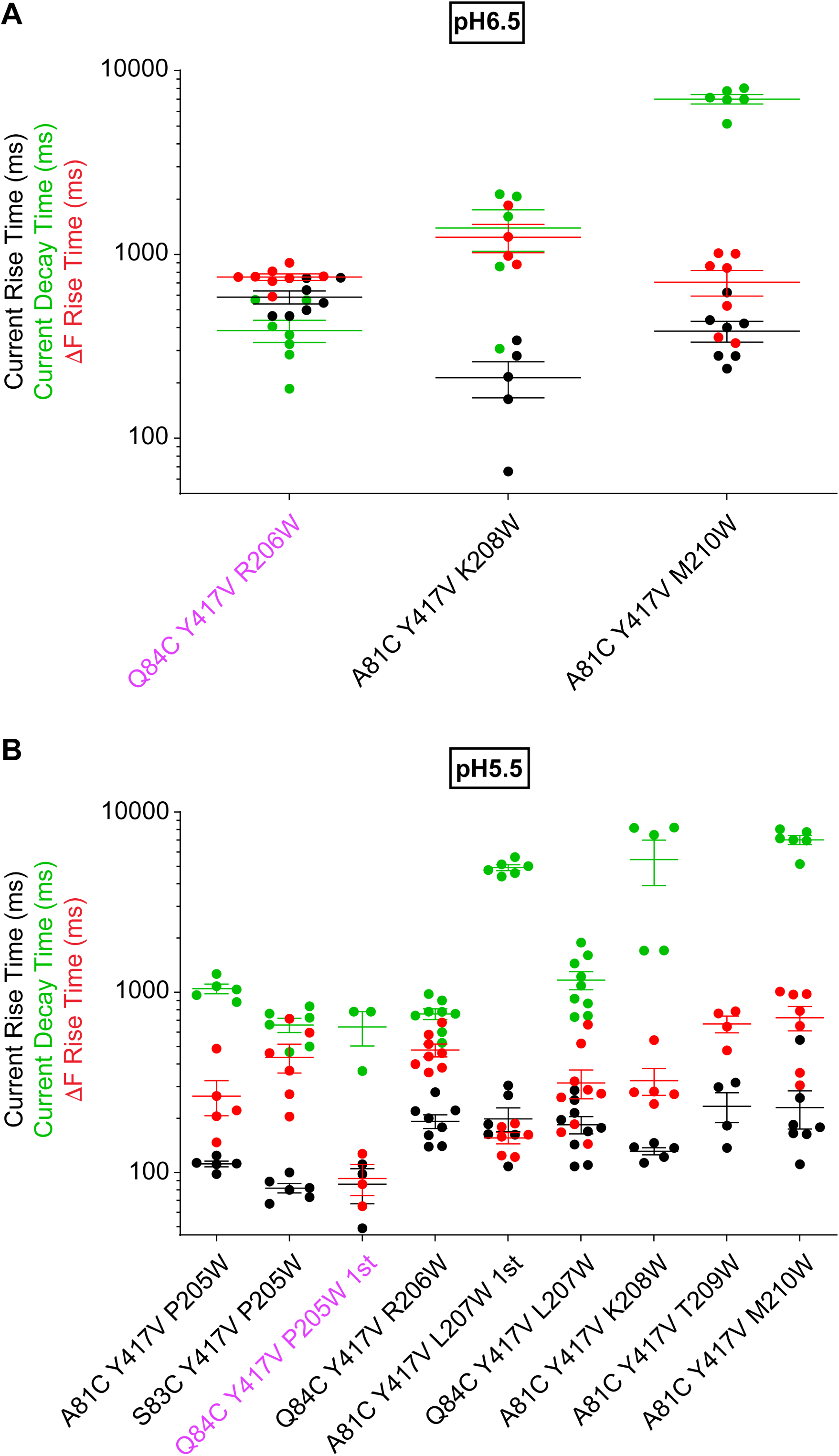
Slow ΔF kinetics of intrasubunit palm triple mutants at pH6.5 and 5.5. Current RT and decay time and ΔF RT obtained at pH6.5 (**A**) or 5.5 (**B**), as indicated, n=3-9. The color of the labels of the mutants indicates that the ΔF onset kinetics are correlated with current appearance (purple; *Materials and Methods*).

**Figure 5-figure supplement 2.**
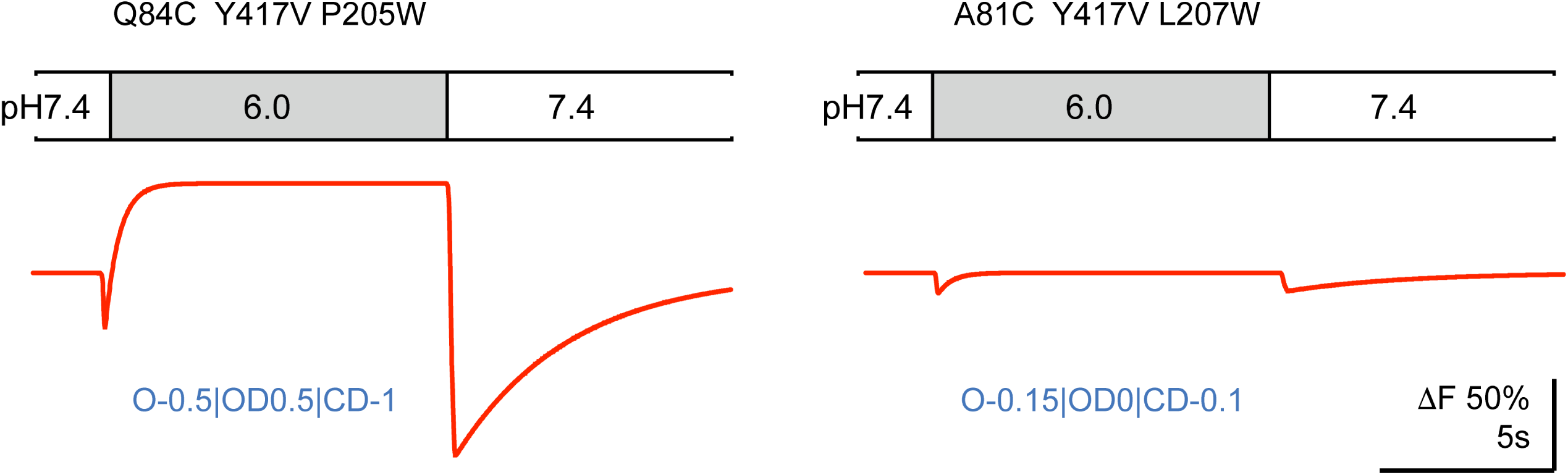
Model-generated ΔF patterns for two mutations of Figure 5. The proportionality factors used for each pattern is indicated below the traces.

**Figure 6-figure supplement 1.**
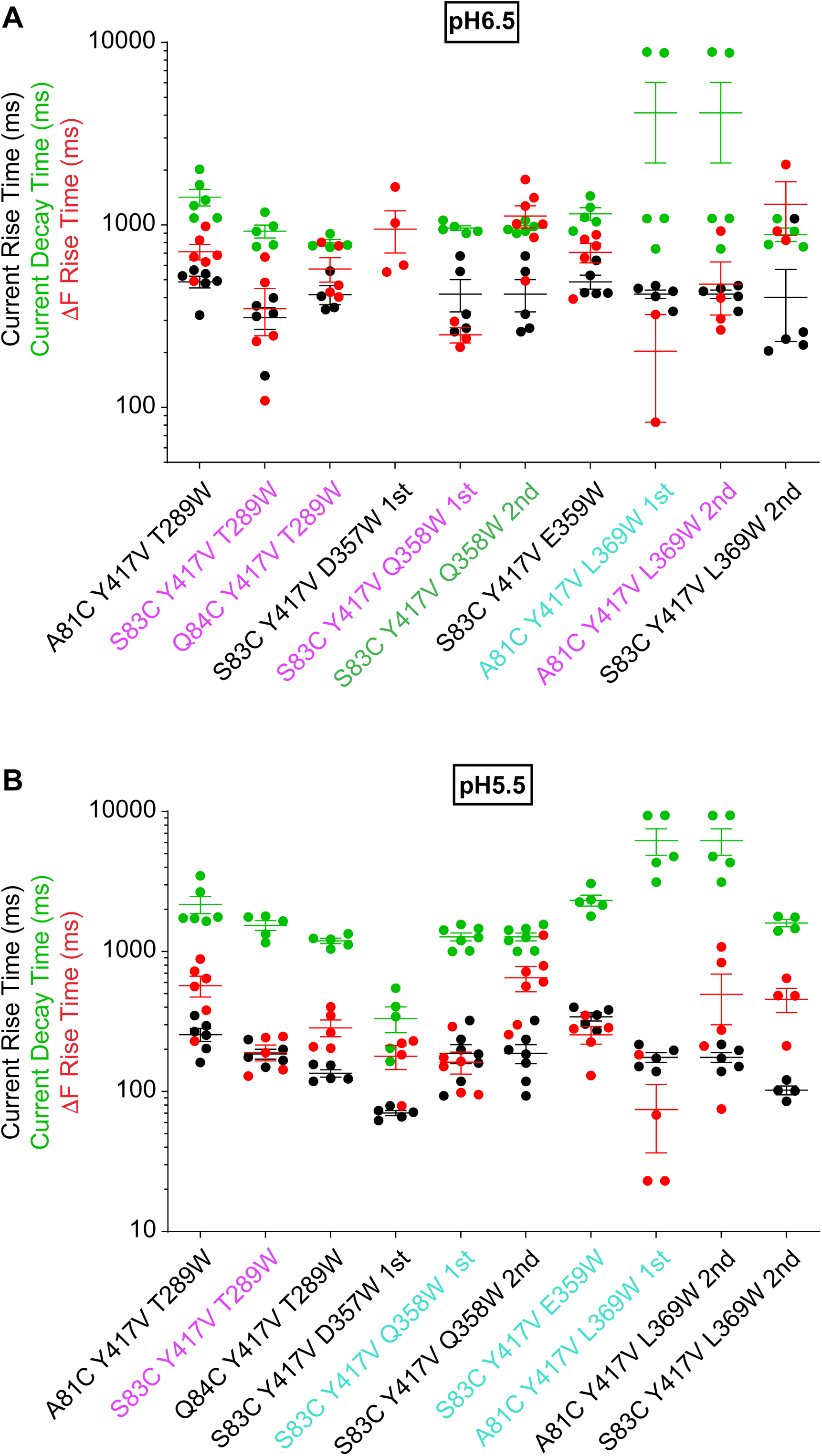
Fast ΔF kinetics of intersubunit palm triple mutants at pH6.5 and 5.5. Current RT and decay time and ΔF RT obtained at pH5.5 (**A**) or 6.5 (**B**), as indicated, n=2-7. The color of the labels of the mutants indicates that the ΔF onset kinetics are correlated with current appearance (purple) or decay (green; *Materials and Methods*) or are faster than current appearance (cyan). At pH6.5, the current of the mutant S83C/Y417V/D357W was too small for fitting of the kinetics, therefore only the kinetics of the ΔF signal are shown.

**Figure 6-figure supplement 2.**
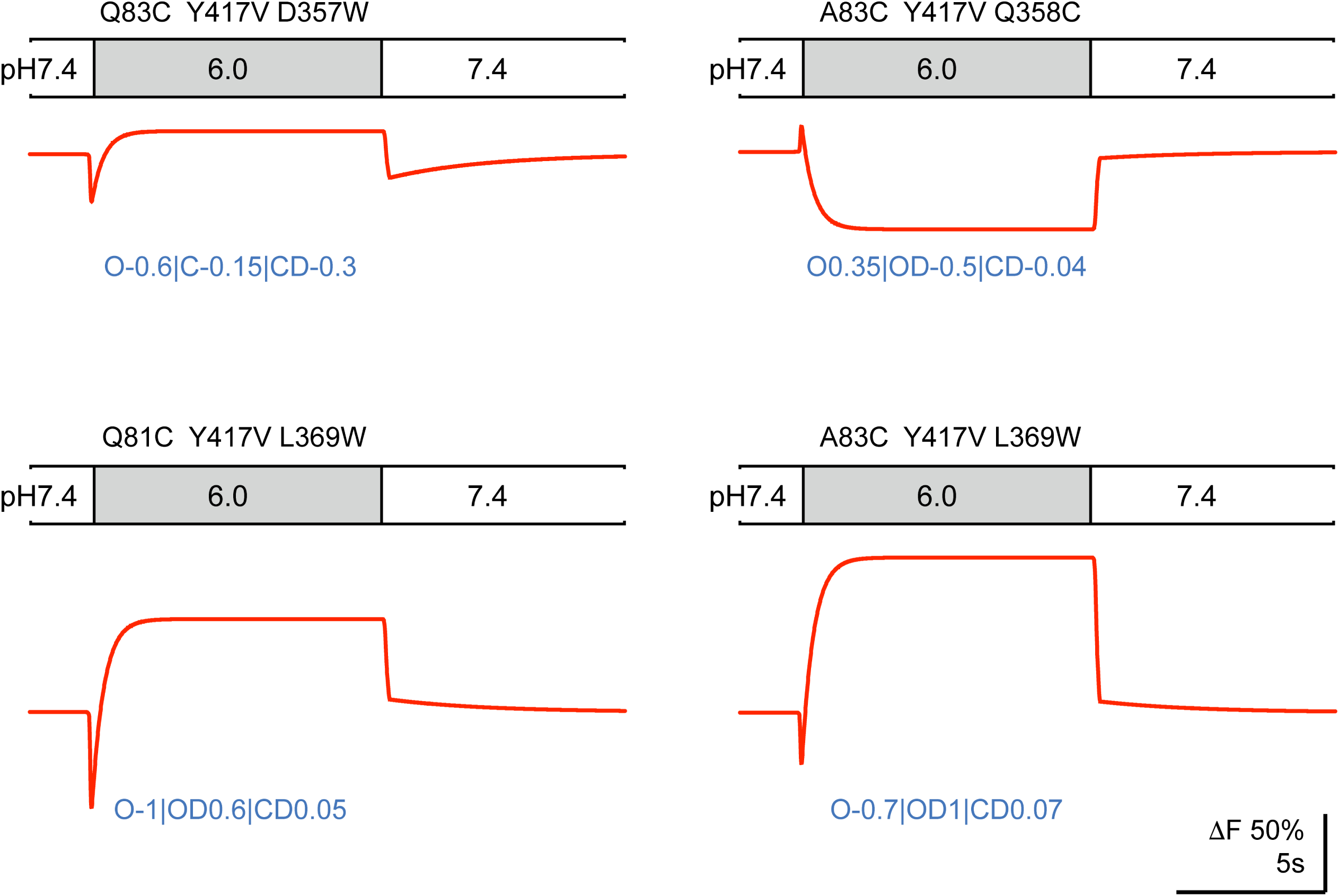
Model-generated ΔF patterns for four mutants of Figure 6. The proportionality factors used for each pattern are indicated below the traces.

**Figure 6-figure supplement 3.**
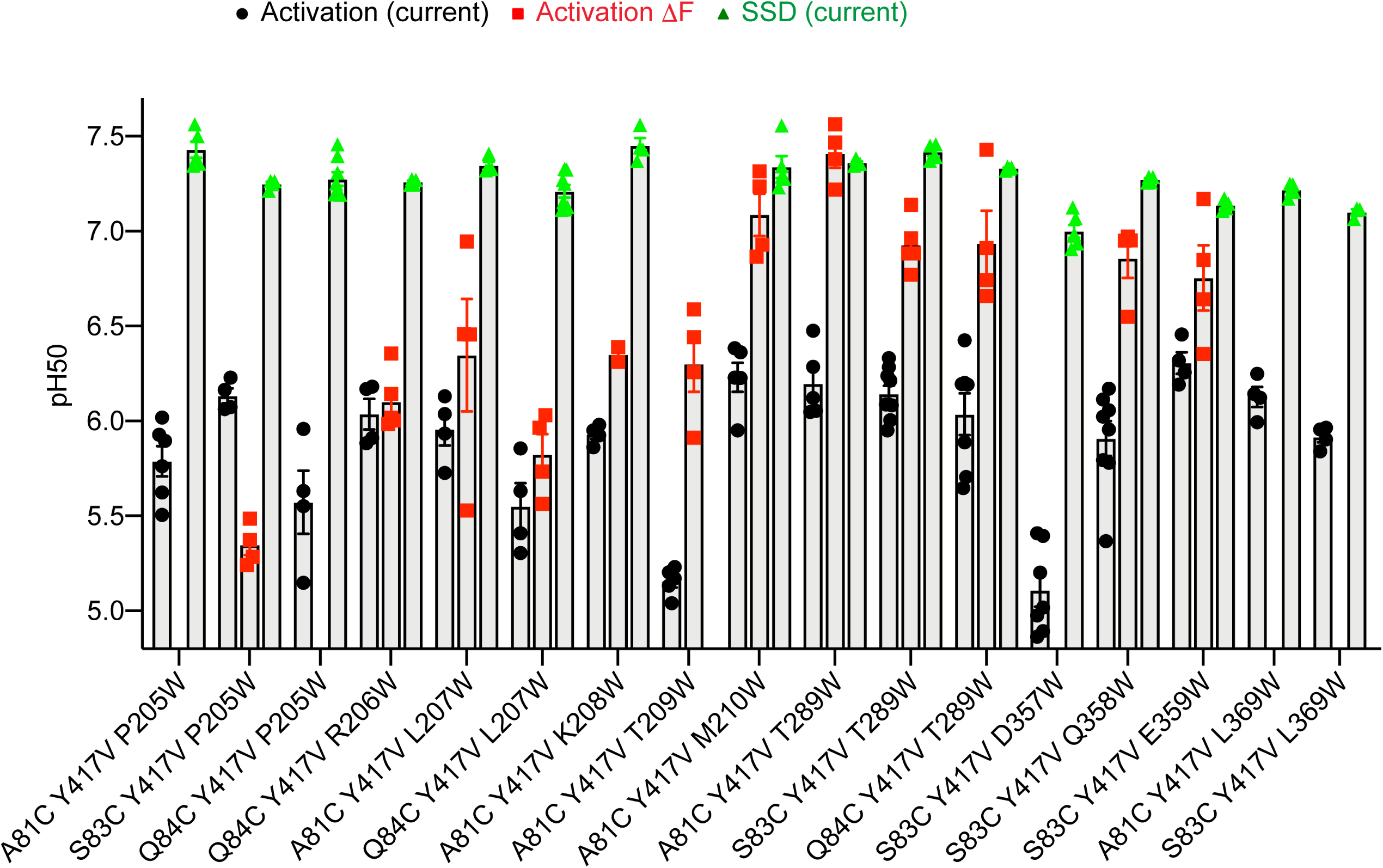
pH dependence of palm mutants. pH_50_ values of current activation (black), current SSD (green) and ΔF amplitude (red) are indicated, n = 2-9. Note that ΔF pH_50_ values are not indicated for all mutants.

**Figure 6-figure supplement 4.**
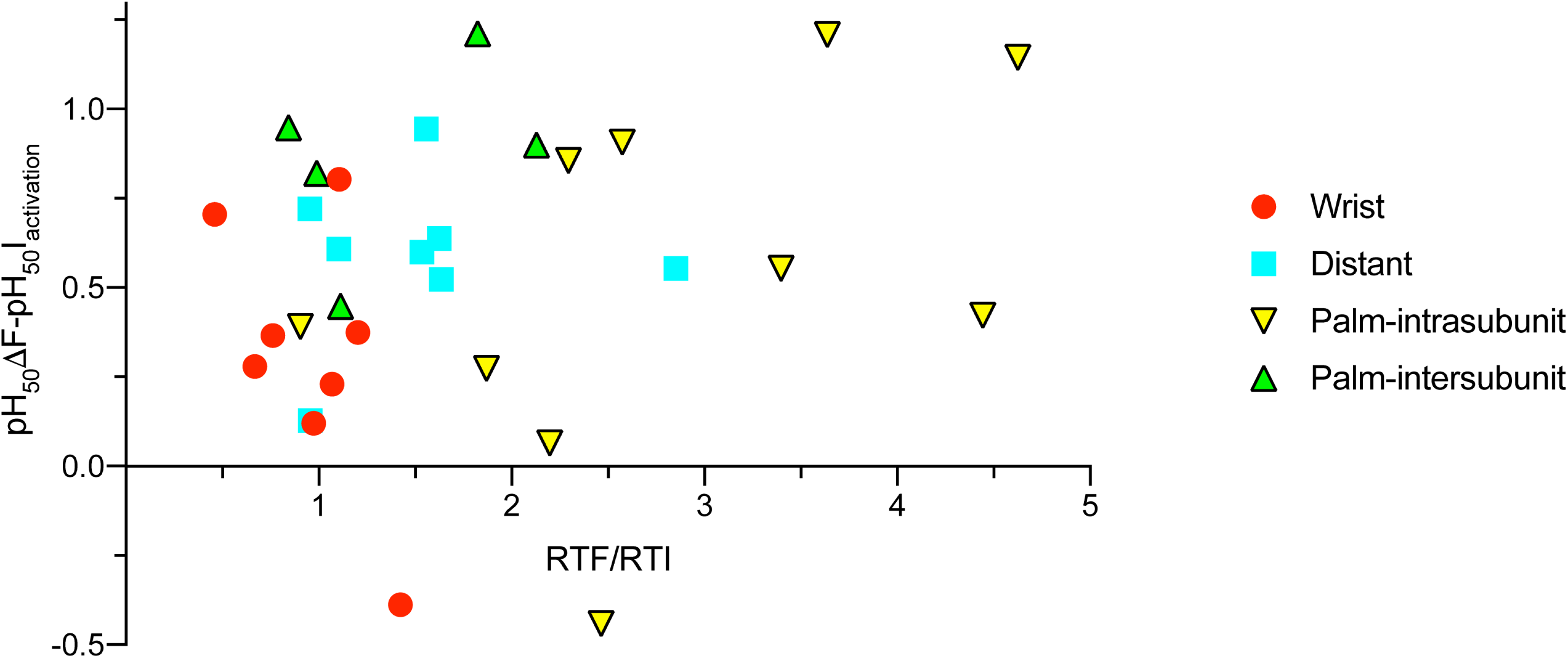
Correlation between the ΔF - current pH_50_ difference and ΔF kinetics. The mean value of the pH_50_(ΔF)-pH_50_(current activation) difference is plotted as a function of the ratio of the rise time of ΔF / rise time of current activation, for the mutants analyzed in this study in different areas of ASIC1a. “Distant” refers to the mutants of the finger, acidic pocket and knuckle, presented in Figure 1.

**Figure 7-figure supplement 1.**
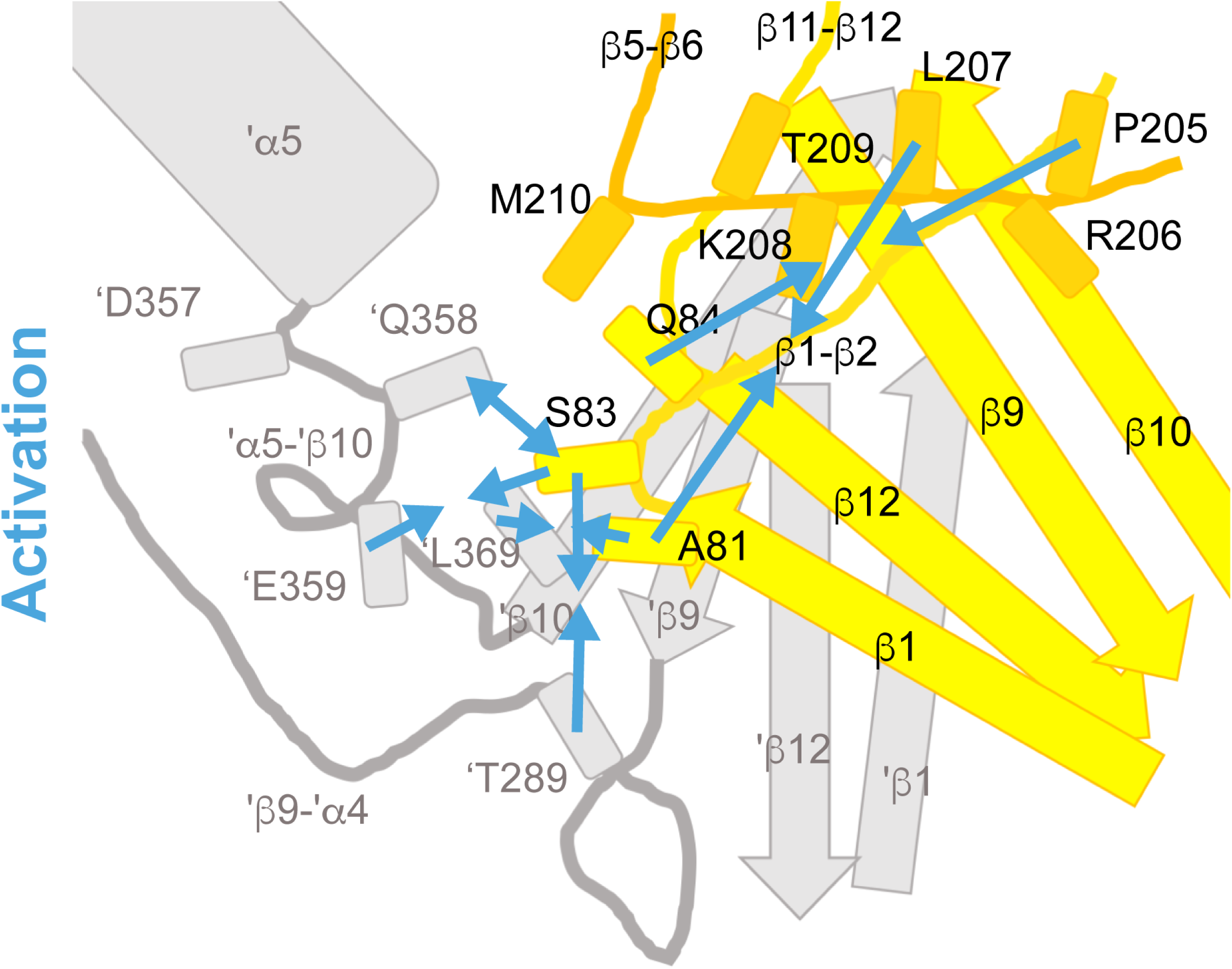
Cartoon of conformational changes during opening predicted by VCF. Cartoon indicating the conformational changes in the palm and palm-thumb loops predicted by VCF to occur during ASIC activation, before validation by SMD. The lower palm domain of one subunit is shown in yellow, structural elements of a neighboring subunit are shown in grey. Arrows that point toward each other indicate an approaching between residues, while arrows pointing away from each other indicate an increase in distance.

**Figure 8-figure supplement 1.**
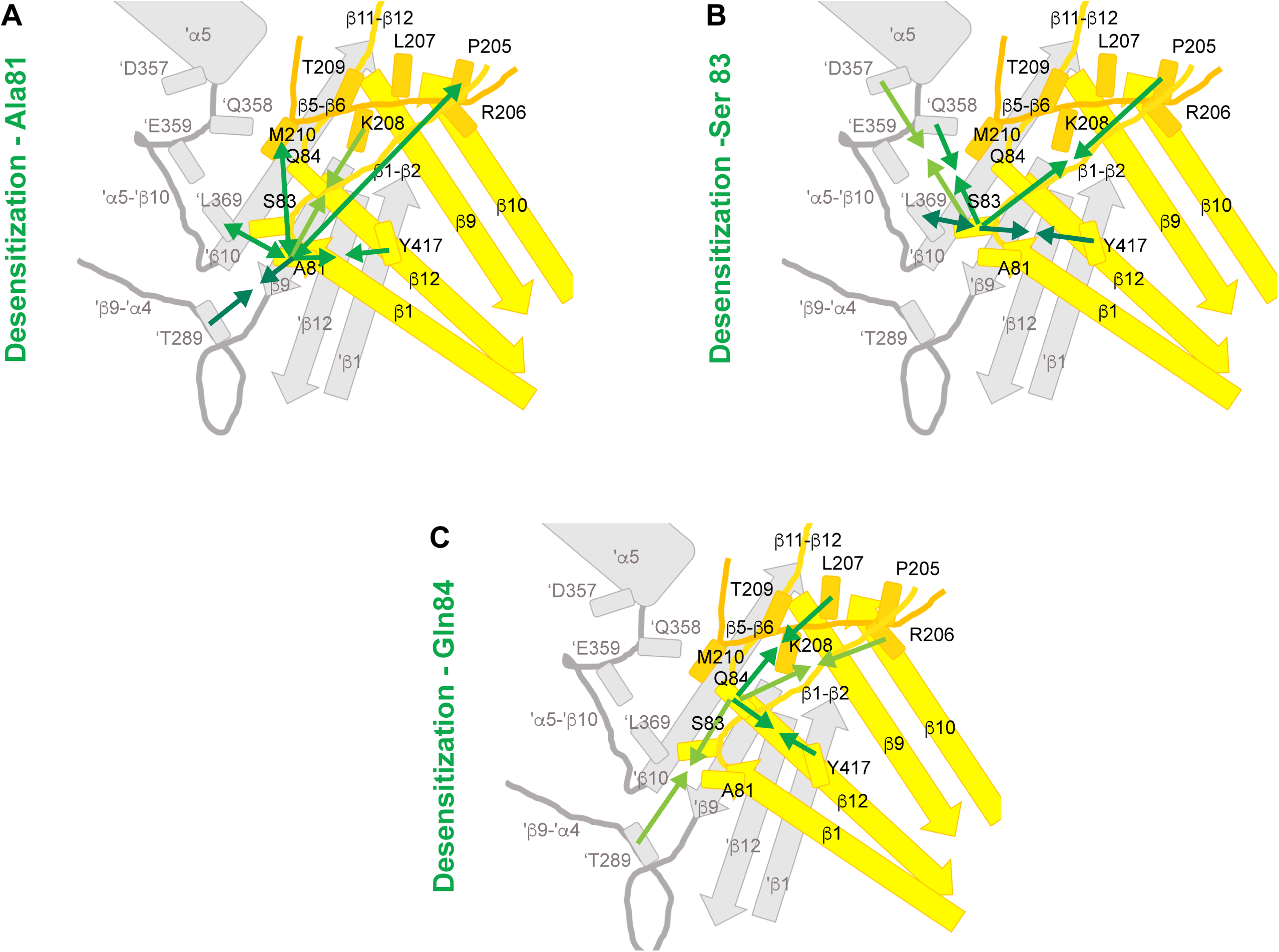
Cartoons of conformational changes during desensitization predicted by VCF. Cartoons indicated the conformational changes in the palm and palm-thumb loops predicted by VCF to occur during ASIC desensitization, before validation by SMD. Predicted distance changes relative to Ala81 (**A**), Ser83 (**B**) and Gln84 (**C**) are shown. The lower palm domain of one subunit is shown in yellow, structural elements of a neighboring subunit are shown in grey. Arrows that point toward each other indicate an approaching between residues, while arrows pointing away from each other indicate an increase in distance.

## References

Alijevic, O., Bignucolo, O., Hichri, E., Peng, Z., Kucera, J. P. & Kellenberger, S. 2020. Slowing of the Time Course of Acidification Decreases the Acid-Sensing Ion Channel 1a Current Amplitude and Modulates Action Potential Firing in Neurons. Front Cell Neurosci, 14, 41.

Baconguis, I., Bohlen, C. J., Goehring, A., Julius, D. & Gouaux, E. 2014. X-ray structure of Acid-sensing ion channel 1-snake toxin complex reveals open state of a Na^+^-selective channel. Cell, 156, 717–29.

Baconguis, I. & Gouaux, E. 2012. Structural plasticity and dynamic selectivity of acid-sensing ion channel-spider toxin complexes. Nature, 489, 400–405.

Bassler, E. L., Ngo-Anh, T. J., Geisler, H. S., Ruppersberg, J. P. & Grunder, S. 2001. Molecular and functional characterization of acid-sensing ion channel (ASIC) 1b. J. Biol. Chem., 276, 33782–33787.

Biasini, M., Bienert, S., Waterhouse, A., Arnold, K., Studer, G., Schmidt, T., Kiefer, F., Gallo Cassarino, T., Bertoni, M., Bordoli, L. & Schwede, T. 2014. SWISS-MODEL: modelling protein tertiary and quaternary structure using evolutionary information. Nucleic Acids Res, 42, W252–8.

Bignucolo, O. & Berneche, S. 2020. The Voltage-Dependent Deactivation of the KvAP Channel Involves the Breakage of Its S4 Helix. Front Mol Biosci, 7, 162.

Bignucolo, O., Vullo, S., Ambrosio, N., Gautschi, I. & Kellenberger, S. 2020. Structural and Functional Analysis of Gly212 Mutants Reveals the Importance of Intersubunit Interactions in ASIC1a Channel Function. Front Mol Biosci, 7, 58.

Bonifacio, G., Lelli, C. I. & Kellenberger, S. 2014. Protonation controls ASIC1a activity via coordinated movements in multiple domains. J Gen Physiol, 143, 105–18.

Cha, A. & Bezanilla, F. 1997. Characterizing voltage-dependent conformational changes in the shaker K^+^ channel with fluorescence. Neuron, 19, 1127–1140.

Coric, T., Zhang, P., Todorovic, N. & Canessa, C. M. 2003. The extracellular domain determines the kinetics of desensitization in acid-sensitive ion channel 1. J. Biol. Chem., 278, 45240–45247.

Dahan, D. S., Dibas, M. I., Petersson, E. J., Auyeung, V. C., Chanda, B., Bezanilla, F., Dougherty, D. A. & Lester, H. A. 2004. A fluorophore attached to nicotinic acetylcholine receptor beta M2 detects productive binding of agonist to the alpha delta site. Proc. Natl. Acad. Sci. U S A, 101, 10195–200.

Dawson, R. J., Benz, J., Stohler, P., Tetaz, T., Joseph, C., Huber, S., Schmid, G., Hugin, D., Pflimlin, P., Trube, G., Rudolph, M. G., Hennig, M. & Ruf, A. 2012. Structure of the Acid-sensing ion channel 1 in complex with the gating modifier Psalmotoxin 1. Nat Commun, 3, 936.

Gandhi, C. S. & Olcese, R. 2008. The voltage-clamp fluorometry technique. Methods Mol Biol, 491, 213–31.

Garcia-Anoveros, J., Derfler, B., Nevillegolden, J., Hyman, B. T. & Corey, D. P. 1997. BNaC1 and BNaC2 constitute at new family of human neuronal sodium channels related to degenerins and epithelial sodium channels. Proc. Natl. Acad. Sci. USA, 94, 1459–1464.

Gonzales, E. B., Kawate, T. & Gouaux, E. 2009. Pore architecture and ion sites in acid-sensing ion channels and P2X receptors. Nature, 460, 599–604.

Grunder, S. & Pusch, M. 2015. Biophysical properties of acid-sensing ion channels (ASICs). Neuropharmacology, 94, 9–18.

Gwiazda, K., Bonifacio, G., Vullo, S. & Kellenberger, S. 2015. Extracellular Subunit Interactions Control Transitions between Functional States of Acid-sensing Ion Channel 1a. J Biol Chem, 290, 17956–66.

Humphrey, W., Dalke, A. & Schulten, K. 1996. VMD: visual molecular dynamics. J Mol Graph, 14, 33-8, 27–8.

Jasti, J., Furukawa, H., Gonzales, E. B. & Gouaux, E. 2007. Structure of acid-sensing ion channel 1 at 1.9 A resolution and low pH. Nature, 449, 316–23.

Jing, L., Jiang, Y. Q., Jiang, Q., Wang, B., Chu, X. P. & Zha, X. M. 2011. The interaction between the first transmembrane domain and the thumb of ASIC1a is critical for its N-glycosylation and trafficking. PLoS One, 6, e26909.

Jo, S., Kim, T. & Im, W. 2007. Automated builder and database of protein/membrane complexes for molecular dynamics simulations. PLoS One, 2, e880.

Kellenberger, S. & Schild, L. 2015. International Union of Basic and Clinical Pharmacology. XCI. Structure, Function, and Pharmacology of Acid-Sensing Ion Channels and the Epithelial Na^+^ Channel. Pharmacol Rev, 67, 1–35.

Klauda, J. B., Venable, R. M., Freites, J. A., O’connor, J. W., Tobias, D. J., Mondragon-Ramirez, C., Vorobyov, I., Mackerell, A. D. & Pastor, R. W. 2010. Update of the CHARMM All-Atom Additive Force Field for Lipids: Validation on Six Lipid Types. Journal of Physical Chemistry B, 114, 7830–7843.

Krauson, A. J., Rued, A. C. & Carattino, M. D. 2013. Independent contribution of extracellular proton binding sites to ASIC1a activation. J Biol Chem, 288, 34375–83.

Li, T., Yang, Y. & Canessa, C. M. 2009. Interaction of the aromatics Tyr-72/Trp-288 in the interface of the extracellular and transmembrane domains is essential for proton gating of acid-sensing ion channels. J. Biol. Chem., 284, 4689–94.

Liechti, L. A., Berneche, S., Bargeton, B., Iwaszkiewicz, J., Roy, S., Michielin, O. & Kellenberger, S. 2010. A combined computational and functional approach identifies new residues involved in pH-dependent gating of ASIC1a. J Biol Chem, 285, 16315–29.

Lynagh, T., Mikhaleva, Y., Colding, J. M., Glover, J. C. & Pless, S. A. 2018. Acid-sensing ion channels emerged over 600 Mya and are conserved throughout the deuterostomes. Proc Natl Acad Sci U S A, 115, 8430–8435.

Mackerell, A. D., Bashford, D., Bellott, M., Dunbrack, R. L., Evanseck, J. D., Field, M. J., Fischer, S., Gao, J., Guo, H., Ha, S., Joseph-Mccarthy, D., Kuchnir, L., Kuczera, K., Lau, F. T., Mattos, C., Michnick, S., Ngo, T., Nguyen, D. T., Prodhom, B., Reiher, W. E., Roux, B., Schlenkrich, M., Smith, J. C., Stote, R., Straub, J., Watanabe, M., Wiorkiewicz-Kuczera, J., Yin, D. & Karplus, M. 1998. All-atom empirical potential for molecular modeling and dynamics studies of proteins. J Phys Chem B, 102, 3586–616.

Mackerell, A. D., Feig, M. & Brooks, C. L. 2004. Extending the treatment of backbone energetics in protein force fields: Limitations of gas-phase quantum mechanics in reproducing protein conformational distributions in molecular dynamics simulations. Journal of Computational Chemistry, 25, 1400–1415.

Mannuzzu, L. M., Moronne, M. M. & Isacoff, E. Y. 1996. Direct physical measure of conformational rearrangement underlying potassium channel gating. Science, 271, 213–6.

Mansoor, S. E., Mchaourab, H. S. & Farrens, D. L. 2002. Mapping proximity within proteins using fluorescence spectroscopy. A study of T4 lysozyme showing that tryptophan residues quench bimane fluorescence. Biochemistry, 41, 2475-84.

Pantazis, A. & Olcese, R. 2012. Relative transmembrane segment rearrangements during BK channel activation resolved by structurally assigned fluorophore-quencher pairing. J. Gen. Physiol., 140, 207–18.

Pantazis, A., Westerberg, K., Althoff, T., Abramson, J. & Olcese, R. 2018. Harnessing photoinduced electron transfer to optically determine protein sub-nanoscale atomic distances. Nat Commun, 9, 4738.

Paukert, M., Chen, X., Polleichtner, G., Schindelin, H. & Grunder, S. 2008. Candidate amino acids involved in H^+^ gating of acid-sensing ion channel 1a. J. Biol. Chem., 283, 572–81.

Pettersen, E. F., Goddard, T. D., Huang, C. C., Couch, G. S., Greenblatt, D. M., Meng, E. C. & Ferrin, T. E. 2004. UCSF Chimera--a visualization system for exploratory research and analysis. J. Comput. Chem., 25, 1605–12.

Pless, S. A. & Lynch, J. W. 2009. Ligand-specific conformational changes in the alpha1 glycine receptor ligand-binding domain. J. Biol. Chem., 284, 15847–56.

Rook, M. L., Williamson, A., Lueck, J. D., Musgaard, M. & Maclean, D. M. 2020. beta11-12 linker isomerization governs acid-sensing ion channel desensitization and recovery. Elife, 9.

Roy, S., Boiteux, C., Alijevic, O., Liang, C., Berneche, S. & Kellenberger, S. 2013. Molecular determinants of desensitization in an ENaC/degenerin channel. FASEB J, 27, 5034–45.

Schuhmacher, L. N., Srivats, S. & Smith, E. S. 2015. Structural domains underlying the activation of acid-sensing ion channel 2a. Mol Pharmacol, 87, 561–71.

Springauf, A., Bresenitz, P. & Grunder, S. 2011. The interaction between two extracellular linker regions controls sustained opening of acid-sensing ion channel 1. J Biol Chem, 286, 24374–84.

Sun, D., Liu, S., Li, S., Zhang, M., Yang, F., Wen, M., Shi, P., Wang, T., Pan, M., Chang, S., Zhang, X., Zhang, L., Tian, C. & Liu, L. 2020. Structural Insights into Human Acid-1 sensing Ion Channel 1a Inhibition by Snake Toxin Mambalgin1. e-life, in press.

Sutherland, S. P., Benson, C. J., Adelman, J. P. & Mccleskey, E. W. 2001. Acid-sensing ion channel 3 matches the acid-gated current in cardiac ischemia-sensing neurons. Proc. Natl. Acad. Sci. USA, 98, 711–716.

Vaithia, A., Vullo, S., Peng, Z., Alijevic, O. & Kellenberger, S. 2019. Accelerated Current Decay Kinetics of a Rare Human Acid-Sensing ion Channel 1a Variant That Is Used in Many Studies as Wild Type. Front Mol Neurosci, 12, 133.

Vullo, S., Bonifacio, G., Roy, S., Johner, N., Berneche, S. & Kellenberger, S. 2017. Conformational dynamics and role of the acidic pocket in ASIC pH-dependent gating. Proc Natl Acad Sci U S A, 114, 3768–3773.

Waldmann, R., Champigny, G., Bassilana, F., Heurteaux, C. & Lazdunski, M. 1997. A proton-gated cation channel involved in acid-sensing. Nature, 386, 173–7.

Wemmie, J. A., Taugher, R. J. & Kreple, C. J. 2013. Acid-sensing ion channels in pain and disease. Nat Rev Neurosci, 14, 461–71.

Wu, Y., Chen, Z. & Canessa, C. M. 2019. A valve-like mechanism controls desensitization of functional mammalian isoforms of acid-sensing ion channels. Elife, 8.

Yandell, B. S. 1997. Practical Data Analysis for Designed Experiments, Chapman and Hall.

Yang, H., Yu, Y., Li, W. G., Yu, F., Cao, H., Xu, T. L. & Jiang, H. 2009. Inherent dynamics of the acid-sensing ion channel 1 correlates with the gating mechanism. PLoS Biol, 7, e1000151.

Yang, L. & Palmer, L. G. 2014. Ion conduction and selectivity in acid-sensing ion channel 1. J Gen Physiol, 144, 245–255.

Yoder, N. & Gouaux, E. 2018. Divalent cation and chloride ion sites of chicken acid sensing ion channel 1a elucidated by x-ray crystallography. PLoS One, 13, e0202134.

Yoder, N. & Gouaux, E. 2020. The His-Gly motif of acid-sensing ion channels resides in a reentrant ‘loop’ implicated in gating and ion selectivity. Elife, 9.

Yoder, N., Yoshioka, C. & Gouaux, E. 2018. Gating mechanisms of acid-sensing ion channels. Nature, 555, 397–401.

